# Factorizing polygenic epistasis improves prediction and uncovers biological pathways in complex traits

**DOI:** 10.1101/2022.11.29.518075

**Authors:** David Tang, Jerome Freudenberg, Andy Dahl

## Abstract

Epistasis is central in many domains of biology, but it has not yet proven useful for complex traits. This is partly because complex trait epistasis involves polygenic interactions that are poorly captured in current models. To address this gap, we develop a new model called Epistasis Factor Analysis (EFA). EFA assumes that polygenic epistasis can be factorized into interactions between a few Epistasis Factors (EFs), which represent latent polygenic components of the observed complex trait. The statistical goals of EFA are to improve polygenic prediction and to increase power to detect epistasis, while the biological goal is to unravel genetic effects into more-homogeneous units. We mathematically characterize EFA and use simulations to show that EFA outperforms current epistasis models when its assumptions approximately hold. Applied to predicting yeast growth rates, EFA outperforms the additive model for several traits with large epistasis heritability and uniformly outperforms the standard epistasis model. We replicate these prediction improvements in a second dataset. We then apply EFA to four previously-characterized traits in the UK Biobank and find statistically significant epistasis in all four, including two that are robust to scale transformation. Moreover, we find that the inferred EFs partly recover pre-defined biological pathways for two of the traits. Our results demonstrate that more realistic models can identify biologically and statistically meaningful epistasis in complex traits, indicating that epistasis has potential for precision medicine and characterizing the biology underlying GWAS results.

## Introduction

Epistasis refers to interactions between genetic effects on a trait. Epistasis is central in many domains of biology, including rare human disorders [1–3], protein evolution [4, 5], natural selection [6, 7], and functional genomics [8]. In model systems, statistical models of epistasis can be useful for characterizing genetic architecture [9–16], improving genomic selection [17, 18], and unbiasedly screening for unknown genomic mechanisms [19–22]. In striking contrast with the others domains, it remains debated if epistasis matters in complex traits [23–25]. This is primarily because complex trait biology is poorly understood, which severely limits our ability to study epistatic interactions between causal genetic mechanisms.

Unbiased genome-wide statistical tests for epistasis could quantify and characterize the nature of epistasis in complex traits. However, modelling epistasis in complex traits is challenging because they are affected by a large number of genetic variants, i.e., they are polygenic (**Figure 1a**). Although studies have identified some specific examples of epistasis in complex traits [26–28], they have not explained significant missing heritability [29, 30]. We hypothesize this is partly due to shortcomings in current models of complex trait epistasis. Specifically, prior studies have assumed that each epistatic interaction effect is completely random–independent of all other additive effects and interaction effects (**Figure 1b**, [31–33]). This model is mathematically simple but biologically unrealistic and statistically under-powered.

**Figure 1.**
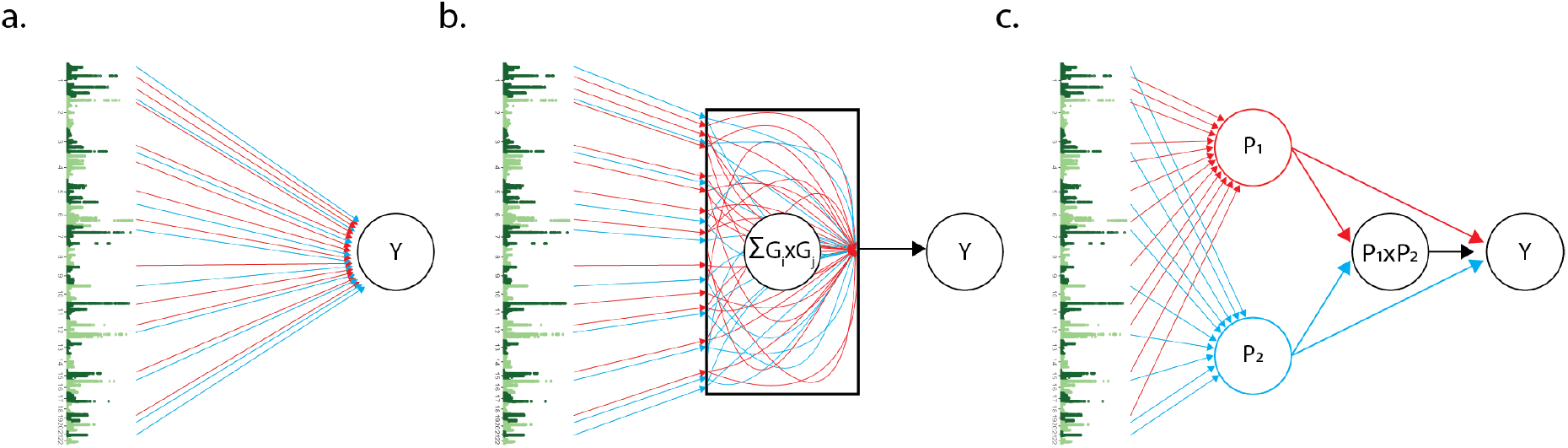
Three genetic architectures consistent with a typical GWAS. (a) Additive: Each SNP has an independent effect on the phenotype, *Y*. (b) Uncoordinated: Each SNP interacts randomly with all other SNPs. (c) Coordinated: SNPs can be grouped into two interacting factors. EFA uses the P1xP2 interaction to (1) improve statistical power to detect epistasis and (2) sort GWAS loci into factors enriched in distinct biological pathways.

Here, we develop a new complex trait epistasis model called Epistasis Factor Analysis (EFA). EFA assumes a “coordinated” form of epistasis where SNP-SNP interactions are structured by interactions between a few latent components, which we call Epistasis Factors (EFs) (**Figure 1c**, [33, 34]). EFA simultaneously partitions GWAS effects into latent EFs and estimates the interactions between the EFs. The inferred EFs can be validated with known genomic annotations and used to discover novel biological pathways underlying GWAS results. A key feature of EFA is leveraging additive effects to improve estimates of epistasis effects, which is important because additive effects are usually much stronger in complex traits. We illustrate this advantage of EFA over standard models in simulations where heritability is mostly additive. We also show EFA improves prediction over the additive model for traits with modest epistasis heritability in two multitrait yeast datasets. We apply EFA to the four UK Biobank traits characterized by [35] and find significant epistasis in all four, including two that are robust to phenotype scale. Finally, we find that the inferred EFs partly recover predefined biological pathways. Our results show that structured models of polygenic epistasis can improve statistical performance and biological understanding in complex traits.

## Epistasis Factor Analysis

Nearly all complex trait studies assume the additive model, and it remains debated whether epistasis matters in complex traits [24, 25]. However, this debate has largely been predicated on an “uncoordinated” model of epistasis that assumes the epistasis effects are completely random–independent of both additive effects and other epistasis effects [30, 31, 36]. Uncoordinated epistasis is not interpretable, realistic, or statistically supported [30]. In contrast, we recently proposed a more realistic “coordinated” model of epistasis where epistasis and additive effects are structured by latent factors (**Figure 1c**, [33]). Coordination is motivated by known forms of epistasis in simpler traits, such as genetic modifiers of a Mendelian gene [2].

Here, we introduce a new coordinated epistasis model to identify these latent factors called Epistasis Factor Analysis (EFA). EFA makes two key assumptions on the nature of epistasis in complex traits:

A1: SNP effects are mediated through a few distinct polygenic factors

A2: These polygenic factors interact

A1 and A2 are biologically motivated. For example, the factors in A1 could reflect distinct causal tissues [37, 38], core genes [39, 40], or heritable exposures like smoking [41]. In accordance with A2, causal tissues might signal each other, core genes might encode proteins that physically interact, or an exposure’s effect may depend on underlying genetic risk.

The key idea in EFA is to share information between additive effects and epistasis effects. On one hand, leveraging additive signals greatly improves power to detect epistasis. On the other hand, leveraging epistasis helps unbiasedly partition SNPs into distinct factors, which is impossible in the additive model without external data or biological annotations.

EFA applies to a quantitative trait measured on *N* individuals, *y*, and a matrix of genotypes measured on *N* individuals at *M* loci, *G*. The EFA model is:

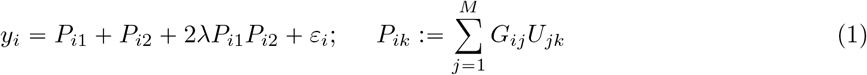

where *P*_*ik*_ is the level of EF *k* for sample *i, U*_*jk*_ is the weight of SNP *j* on EF *k*, and *λ* is the interaction between EFs. We call this Epistasis Factor Analysis because it implicitly makes a low-rank factorization to the genome-wide epistasis matrix (**Methods**). We generalize to *K* > 2 EFs in the **Methods**, and we theoretically characterize EFA in the **Supplement**. Our full model also allows quadratic effects of individual EFs, but we conservatively exclude this term in our main analyses because our goal is to find interactions across biologically distinct pathways.

In practice, *G* will typically contain dozens of SNPs that have been pre-screened based on their additive effects. We fit *U* and *λ* as fixed effects using maximum likelihood (**Methods**). We develop an efficient block coordinate descent algorithm in the **Supplement**. Using linear algebra techniques, we reduce the computational complexity of EFA to the same cost as ordinary linear regression.

### EFA recovers true pathways and improves prediction in simulations

We performed simulations to characterize EFA as a function of additive heritability 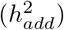, epistasis heritability 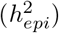, and the fraction of epistasis heritability due to coordinated vs uncoordinated effects (**Methods**). We center and scale genotypes to remove the ambiguity between variance “attributable to” vs variance “caused by” additive effects (in practice, these effects are partly collinear, which can cause confusion [24, 25]). We define broad-sense heritability as 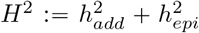, and always use *H*^2^ = 0.5. We compared EFA to uncoordinated epistasis estimates fit either by fixed or random effects (**Methods**). We evaluated estimation accuracy for the pairwise SNP-SNP interactions, *ω*_*jj*_′ (see (3) in the **Methods**), and for the EFs, *U*_*jk*_. EFA directly fits EFs and implicitly fits *ω* (**Methods**). Uncoordinated methods directly fit *ω*, and we derive uncoordinated EFs *post hoc* using PCs of the estimated *ω* matrix (**Methods**).

We first simulated from the EFA model in (1) with *N* = 1, 000 samples and *M* = 20 SNPs. As expected, EFA outperforms uncoordinated estimates of *ω* across the range of 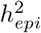 (**Figure 2a**). EFA’s gain is greatest when 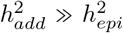, illustrating how EFA’s epistasis estimates borrow strength from additive effects. Analogously, EFA improves additive effect estimates over standard additive models when 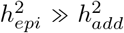 (**Supplementary Figure S1**), though this scenario is not likely in practice. EFA also substantially outperforms uncoordinated estimates of EFs, which is expected because EFA specifically targets these low-dimensional factors (**Figure 2b**). This also shows how EFA leverages additive signals to improve EF estimates: when 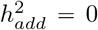, EFA actually underperforms the uncoordinated model, whereas EFA remains accurate even when heritability is 90% additive.

**Figure 2.**
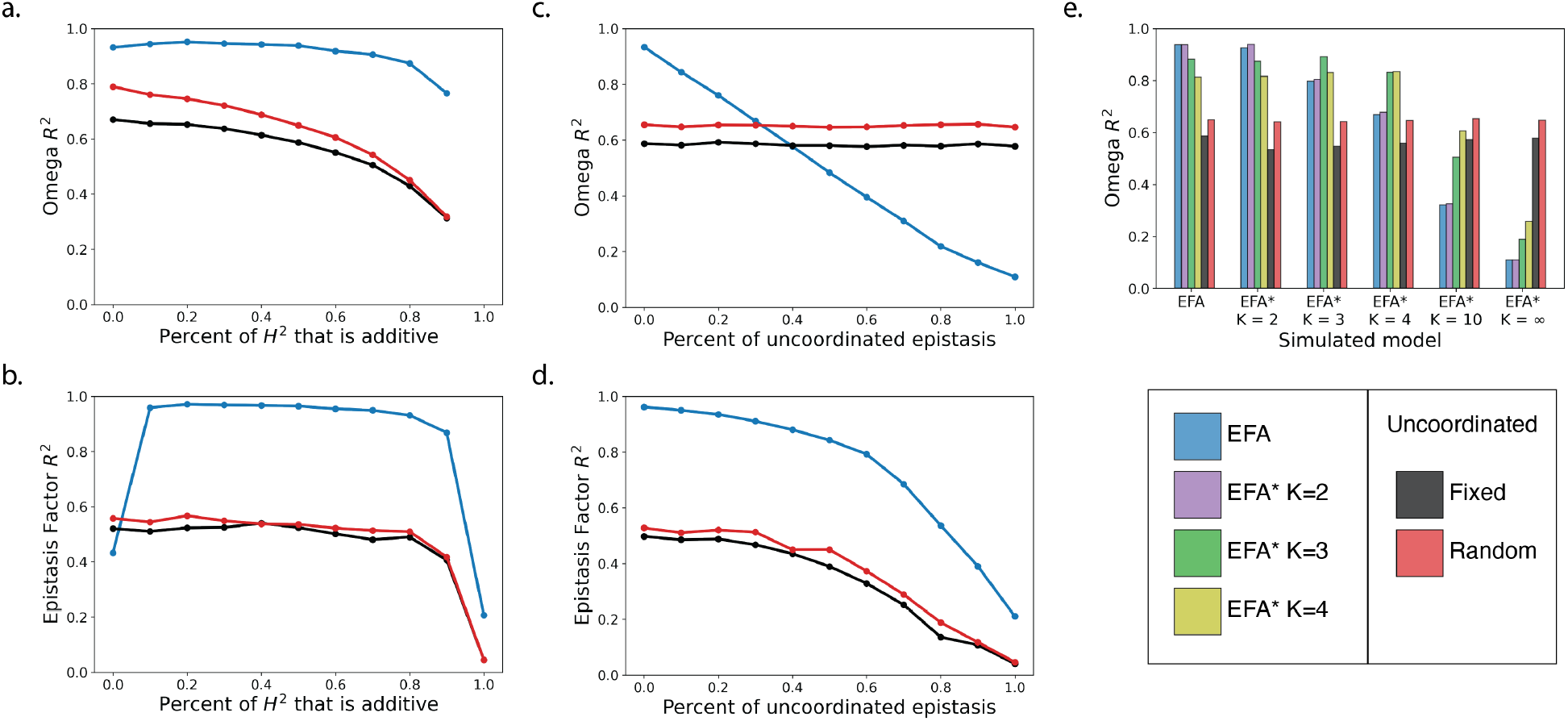
Simulations characterize the accuracy of polygenic epistasis estimates. Accuracy is quantified by the squared correlation between estimated and true pairwise SNP epistasis effects, *ω* (a,c,e) or between estimated and true EFs, *U* (b,d). Simulations under the EFA model vary the fraction of heritability due to additive vs epistasis effects (a,b). Partial coordination is simulated by combining epistasis effects from the EFA model with uncoordinated epistasis effects (c,d). Other forms of coordination are simulated using the EFA^*^ model with *K* ⩾ 2 (e), which is a more general version of EFA (**Methods**). In (b,d), the EF *R*^2^ is nonzero even without any coordinated epistasis because the EFs still capture the additive effects.

We then fix 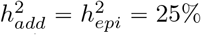 and vary the level of coordination by adding in uncoordinated epistasis effects (**Methods**). As expected, EFA’s performance decays as coordination weakens, while the uncoordinated estimates remain stable (**Figure 2c**). Nonetheless, EFA always outperforms uncoordinated estimates of the EFs because EFA specifically targets the coordinated component of epistasis (**Figure 2d**).

We next simulated and fit *K* > 2 EFs using a generalization of EFA called EFA^*^ (**Methods**). As expected, *ω* estimates were more accurate when fitting EFA^*^ with the true value of *K* (**Figure 2e**), though the base EFA model was robust to modest misspecification of *K*. However, the uncoordinated model performed better as *K* grows. This is expected, as the *K* =∞ case is equivalent to uncoordinated epistasis [33]; intuitively, the independent pathways become individually negligible and wash out. We did not consider EF estimation accuracy because EFs are not identified for *K* > 2 (**Supplement**).

We next varied the number of SNPs, *M*, and sample size, *N* (**Supplementary Figure S2**). Across all *N* and *M*, EFA always outperforms uncoordinated estimates. Performance for all methods decays as *M* grows because each individual SNP effect is smaller. This illustrates why modelling epistasis in complex traits is difficult. Likewise, all methods improve as *N* grows.

Finally, we asked if EFA can improve phenotype prediction. We first simulated from EFA’s model and found that EFA performed well, with prediction *R*^2^ near the theoretical limit of *H*^2^ (**Figure 3a**). EFA always outperformed the fixed-effect uncoordinated model, and EFA outperformed the random effect uncoordinated model except when epistasis was absent. (We derive the predictions from the uncoordinated random effect model in the **Supplement**.) We then fixed 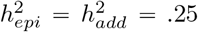 and varied the fraction of epistasis due to coordination, as above. Results were similar to *ω* estimation accuracy: when epistasis is mostly coordinated, EFA is optimal, but EFA performs poorly when epistasis is uncoordinated (**Figure 3b**). Overall, EFA improves estimation of latent pathways in all our tests and improves prediction when epistasis is sufficiently strong and coordinated.

**Figure 3.**
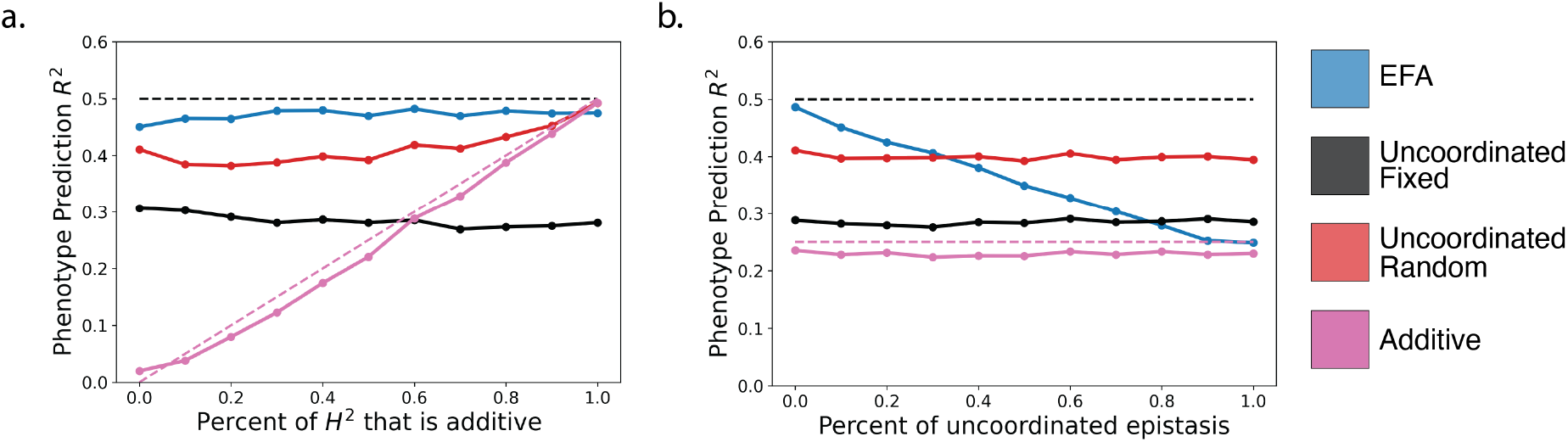
Phenotype prediction accuracy in simulations. (a) varies the fraction of heritability due to additive vs epistasis effects. (b) varies the fraction of epistasis effects due to coordinated pathway-level interactions vs uncoordinated random interactions. Dotted line is the broad-sense heritability, *H*^2^.

### EFA improves prediction in complex yeast traits

We next asked if EFA could improve genetic prediction in complex traits. We studied 46 complex yeast traits measured on 1008 samples, where each trait was growth rate on a different medium [13]. These traits have varying levels of epistasis heritability, as defined by the difference between the phenotypic similarity of clones (*H*^2^) and the additive heritability estimated from genotype data (*h*^2^). We measured prediction accuracy by the squared correlation between predicted and observed phenotypes using 10-fold cross-validation. For each trait, this yielded 30-70 clumped+thresholded SNPs (*r*^2^ < 0.2, **Methods**).

We first compared EFA to the additive model and found that EFA significantly improved prediction accuracy for 6/46 traits (*p* < .05, paired *t*-test, **Methods, Supplementary Table 1**). These 6 traits all had large epistasis heritability (**Figure 4a**). More generally, the difference between 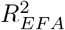 and 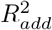 correlated with the epistasis heritability (Spearman’s *ρ* = 0.48, *p* < 0.001). This is 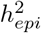 is higher. By comparison, the additive model significantly improved prediction over EFA for 20/46 traits. Finally, EFA predictions were superior to uncoordinated epistasis predictions using fixed effects for all 46 traits (**Figure 4b**).

**Table 1:** EFA finds statistically significant evidence of epistasis in complex traits. Confidence intervals were computed using the 0.025 and 0.975 quantiles of the bootstrap distribution. All *p*-values are two sided.

|  |  | IGF-1 | Urate | Testosterone (Male) | Testosterone (Female) |
| --- | --- | --- | --- | --- | --- |
| Standard | CI | (0.25, 0.60) | (-0.50, -0.27) | (0.39, 0.70) | (1.19, 1.78) |
|  | Empirical p-val | **.002** | **.006** | **.002** | **.002** |
|  | Gaussian p-val | 3.52e-04 | 1.16e-07 | .011 | 2.75e-04 |
| QNorm | CI | (-0.30, 0.37) | (-0.57, -0.36) | (-0.46, 0.42) | (0.63, 1.04) |
|  | Empirical p-val | .570 | **.002** | .826 | **.002** |
|  | Gaussian p-val | .689 | 7.67e-15 | .846 | 3.72e-12 |
| Permuted | CI | (-0.28, 0.32) | (-0.47, 0.00) | (-0.46, 0.41) | (-1.48, 0.75) |
|  | Empirical p-val | .730 | .066 | .618 | .274 |
|  | Gaussian p-val | .749 | .090 | .642 | .984 |

**Figure 4.**
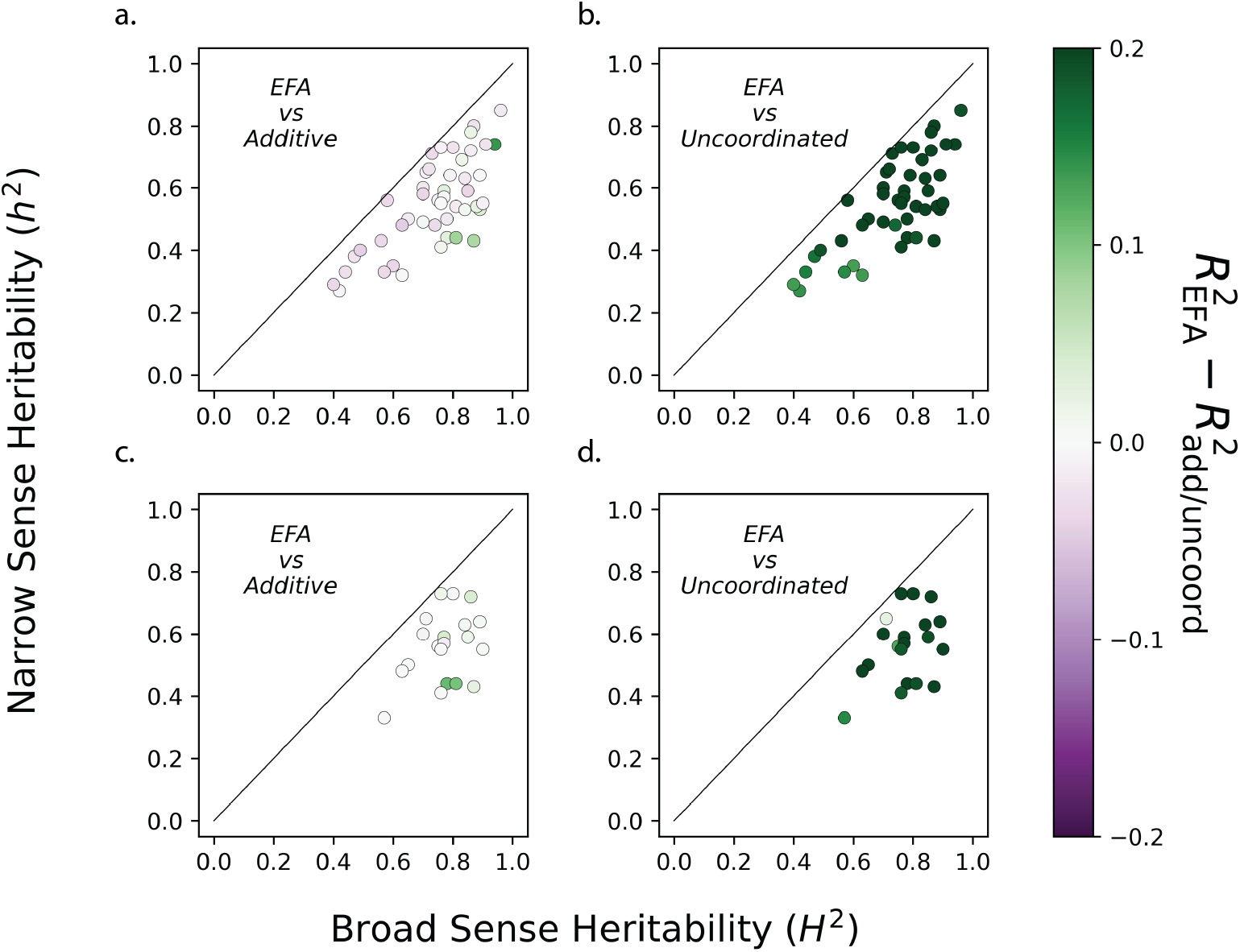
Prediction accuracy in complex yeast traits. Each point is a trait with narrow- and broad-sense heritability estimates from [42]. (a) EFA significantly outperforms the additive model for 6/46 traits from Bloom et al 2013 data. (c) EFA significantly outperforms the additive model for 7/20 traits from Bloom et al 2015 data. (b,d) EFA always outperforms the uncoordinated fixed-effect epistasis model.

Next, we compared EFA to additive and epistasis random effects models. Unlike fixed effect models, including EFA, random effect models can fit the whole genome without LD pruning, which is often a major advantage for prediction in complex traits [18, 43, 44]. Nonetheless, EFA significantly outperforms both additive and uncoordinated epistasis random effect models for 7/46 traits (*p* < .05, **Supplementary Figure S3**). We also compared the two random effect models, and we found that the uncoordinated epistasis model significantly outperformed the additive model for 9/46 traits (*p* < .05). Intriguingly, these 9 traits only overlap 1 of the 7 traits where EFA outperforms the additive random effect model, suggesting that the degree of coordination varies across traits.

Finally, we evaluated replication in a second dataset from the same group with 4390 samples measured on 20 of the above 46 traits [14] (**Supplementary Figure S4**). We replicated the superiority of EFA over the additive model for all 3 traits where EFA beat the additive model in the first dataset (*p* < .05; the other 3/6 traits were not measured in the second dataset). Further, at this larger sample size, we find 4 additional traits where EFA significantly outperforms the additive fixed effect model (**Figure 4c, Supplementary Table 2**). We also replicate that EFA always outperforms the uncoordinated fixed effects model (**Figure 4d**). Overall, EFA improves prediction over the additive model for some complex traits, and the utility of EFA will likely grow with larger sample sizes.

**Table 2:** Epistasis Factors are enriched in predefined biological pathways from [35]. We show pathways that have been assigned > 2 SNPs used in EFA. *p*-values are from a hypergeometric test of the top 10 EF SNPs in each biological pathway relative to the other 90 SNPs.

| Trait | Biological Pathway | # SNPs in EF1 | # SNPs in EF2 | # SNPs total |
| --- | --- | --- | --- | --- |
| IGF-1 | Downstream signaling | 1 ($p = 0.284$) | 0 ($p = 1.000$) | 3 |
| | Growth horm. secr. | 0 ($p = 1.000$) | 1 ($p = 0.361$) | 4 |
| | IGF-1 secretion | 0 ($p = 1.000$) | 1 ($p = 0.361$) | 4 |
| | IGF-1 serum balance | 0 ($p = 1.000$) | 1 ($p = 0.549$) | 7 |
| | Ras signaling | 0 ($p = 1.000$) | 1 ($p = 0.430$) | 5 |
| Urate | Solute transport | 3 ($p = 0.175$) | 5 ($p = 0.007$) ★ | 15 |
| Testosterone (male) | Steroid synthesis & metabolism | 0 ($p = 1.000$) | 1 ($p = 0.478$) | 6 |
| | HPG signaling | 0 ($p = 1.000$) | 0 ($p = 1.000$) | 5 |
| | Serum homeostasis | 0 ($p = 1.000$) | 3 ($p = 0.001$) ★ | 3 |
| Testosterone (female) | Steroid synthesis & metabolism | 1 ($p = 0.638$) | 0 ($p = 1.000$) | 7 |

### EFA detects epistasis and recovers known pathways in human traits

We applied EFA to complex human traits in UK Biobank (UKB). We studied the same four traits that were extensively characterized in [35]: urate, insulin-like growth factor 1 (IGF1), testosterone in males, and testosterone in females. We used the top 100 LD-pruned GWAS hits for each trait (or all 77 for female testosterone, **Methods**).

We first asked if EFA detects statistically significant epistasis using bootstrap. Specifically, we resample individuals to calculate the sampling distribution of *λ*, the factor-level epistasis, which yields empirical *p*-values and confidence intervals (**Methods**). We used simulations to confirm that this bootstrap test is calibrated under the null model where SNPs are additive (**Supplementary Figure S6**). Additionally, we performed realistic simulations with true epistasis and found that EFA’s *λ* estimates are unbiased and that EFA’s bootstrap test is powerful. A minor caveat is that our bootstrap confidence intervals are liberal in the presence of true epistasis, but this does not cause false positive tests of epistasis.

The EFA bootstrap test finds significant epistasis for all four tested traits (all empirical *p* < .05/4, **Figure 5e-h, Table 1**). For IGF-1 and male and female testosterone, *λ* is positive, and for urate, *λ* is negative. Because we use 499 bootstrap replicates, the empirical *p*-values are noisy and lower-bounded by 1/500. If we further assume that our estimate of *λ* is roughly Gaussian under the additive model, we get much lower and more precise *p*-values. Our simulations suggest that this Gaussian approximation is conservative (**Supplementary Figure S7**).

**Figure 5.**
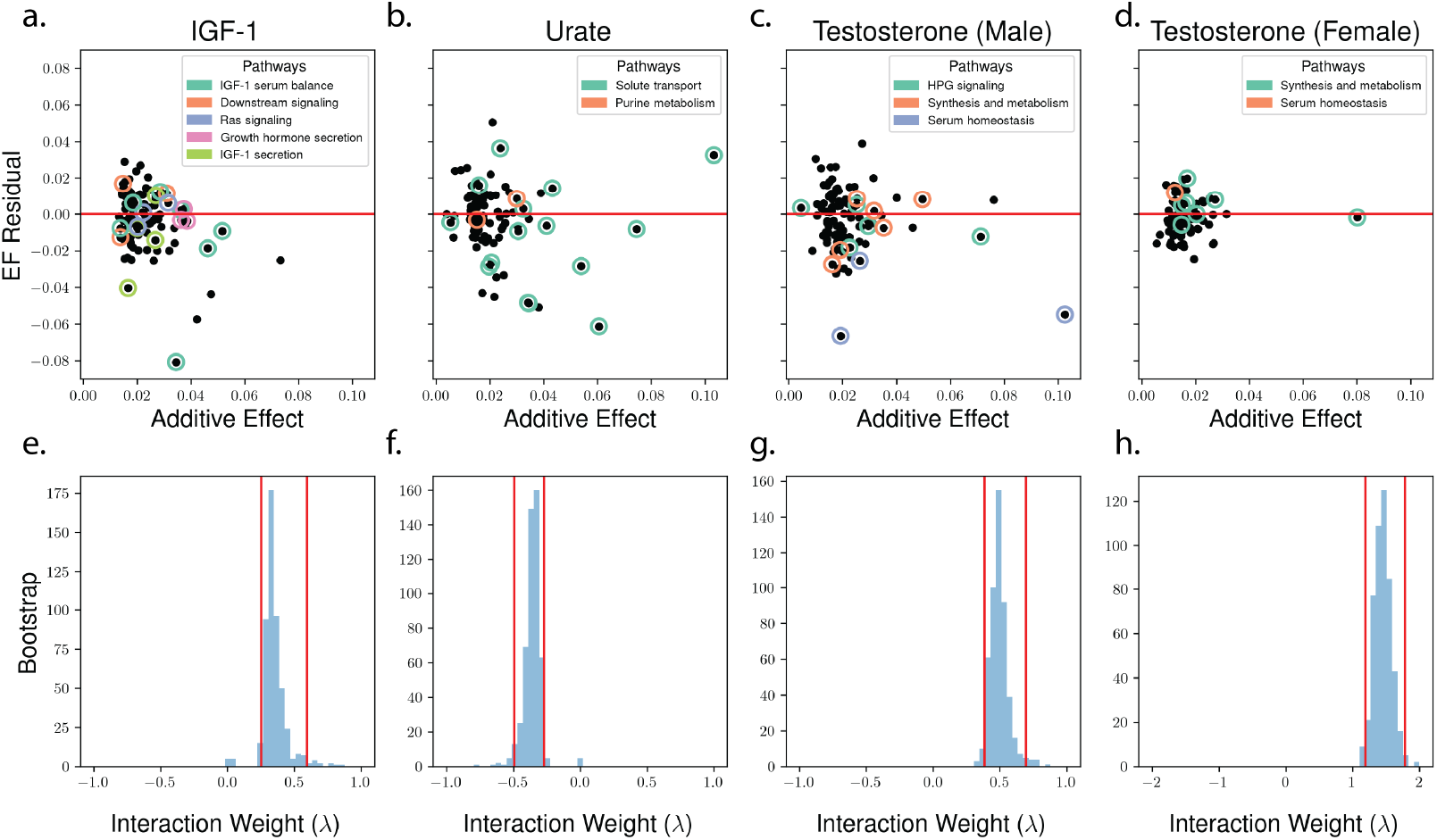
EFA identifies statistically and biologically significant epistasis in 4 complex human traits. (a-d) EFA jointly partitions the additive effects of GWAS-significant SNPs (x-axis) into two EFs (y-axis: difference between the EFs and the additive effects, i.e. *U*_*jk*_ − *β*_*j*_/2). (e-f) Bootstrap replicates show that EFA identifies significant epistasis in all four traits. For visualization, 14 values of *λ* are excluded across all plots (out of a total of 6,000). See **Supplementary Figure S5** for comparable plots on quantile-normalized and permuted phenotypes.

To test robustness under the null, we repeated our analysis after permuting phenotypes. As expected, our bootstrap test was null (**Table 1, Supplementary Figure S5**).

As epistasis can be scale-dependent [45], we conservatively tested robustness by quantile-normalizing phenotypes (**Supplementary Figure S5**). We found that urate and female testosterone remained highly significant (empirical *p* = .002, .002; Gaussian *p* = 7.7 × 10^−15^, 3.7 × 10^−13^). Moreover, the sign and magnitude of *λ* remained stable after this transformation. However, quantile-normalization eliminated the EFA signal for male testosterone and IGF-1, which emphasizes the importance of evaluating scale-dependence in interaction tests. Overall, our results increase confidence that the EFA signals for urate and female testosterone are not scale dependent, while the discrepancies for IGF-1 and male testosterone could either be false positives on the original scale or false negatives on the transformed scale [33].

We next performed simulations to assess EFA’s robustness to dominance and un-genotyped causal variants in LD with genotyped variants. First, we simulated a pure-dominance model without any epistasis between distinct SNPs (**Supplement Section 3.2-3**). Under modest levels of dominance (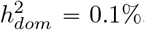, [30, 46]), the bootstrap test is roughly null (**Supplementary Figure S6**). Under strong dominance 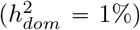, however, the bootstrap test becomes significant (**Supplementary Figure S6**). Both results are expected: dominance is not technically a false positive under EFA (unlike our prior Even-Odd test [33]), but EFA primarily targets inter-SNP epistasis, hence the EFA test has low power if dominance effects are the only form of epistasis.

Second, we simulated an additive model with unobserved causal variants in LD with measured variants. Our simulation meets the necessary condition for ‘phantom’ epistasis in [47], which can cause severe bias in pairwise SNP-SNP epistasis tests (**Supplement**). For EFA, however, we do not observe any bias from ungenotyped variants (**Supplementary Figure S6**). Nonetheless, in principle, other forms of LD could bias EFA estimates. Overall, the structured and polygenic nature of EFA substantially mitigates the LD biases that can plague other epistasis tests [18, 48–51].

We next asked if EFA captures biologically meaningful epistasis by comparing estimated EFs to the known biological pathways defined in [35]. Specifically, we used a hypergeometric test for enrichment of the top 10 SNPs per EF in each predefined biological pathway (relative to the other 90 GWAS SNPs, **Table 2**).

We find that EF2 for male testosterone is enriched in “serum homeostasis” (*p* = 0.0001). This enrichment is driven by SNPs assigned to the genes *SHBG* and *SLCO1B3*, which live on different chromosomes (17 and 12). In urate, EF2 is enriched in “solute transport” (*p* = 0.007, **Table 2**), which is driven solely by a single locus with large effect (*SLC2A9*). This urate signal could either reflect genuine *cis*-*trans* interactions or could merely reflect subtle LD with unmeasured additive effects and/or dominance effects [30]. We confirmed these enrichments are robust if we instead use the top 15 or 20 SNPs per EF (**Supplementary Table 3-6**). The results are also robust if we increase the window size used to assign SNPs to genes (**Supplementary Table 3-6**). Altogether, EFA is capable of recovering biologically-meaningful pathways solely using statistical interactions.

## Discussion

Despite strong evidence for epistasis in many domains of biology, it remains unclear if modelling epistasis adds biological or statistical value in complex traits. We introduced a model to address this question called Epistasis Factor Analysis (EFA). In contrast to existing uncoordinated models of polygenic epistasis, which lack biological motivation, EFA assumes a structured form of polygenic epistasis driven by latent factors (**Figure 1C**). We showed that EFA outperforms additive predictions in several yeast traits. Additionally, we found that EFA can identify epistasis in complex human traits where standard models fail and that the EFs can partly recover known biological pathways without prior information.

We found that EFA can improve prediction in model organisms and simulations, and it is likely that epistasis will eventually improve prediction in natural populations. However, the within-population prediction gain will likely be modest because current additive models partly capture epistasis effects–especially when causal variants have been driven to lower frequencies by natural selection [24, 52]). It is possible that epistasis predictions will improve portability across ancestries, but epistasis has not yet been shown to contribute to this important problem [53, 54]. Therefore, the greater promise of epistasis models in complex human traits is not better statistical prediction, but rather better biological understanding of GWAS results. Our results showing that EFs are enriched in predefined genomic annotations is a step in this direction.

Many polygenic epistasis models have been previously developed. The standard model, which we call “uncoordinated” [33], assumes that all SNP-SNP epistasis effects are independent of each other and of additive effects [31]. While uncoordinated models are mathematically simple and rigorous, they are biologically implausible and statistically under-powered [30]. More recent polygenic epistasis models make parsimonious assumptions to improve power and interpretation. For example, autosome-sex interactions represent a particular form of structured epistasis that is strongly supported in diverse complex traits [55–58] (though these interactions are at least partly due to gender, not sex). As another example, interactions between a single locus and a polygenic background can identify epistasis hubs [59–64]. Finally, BANN models complex traits as epistatic compositions of simpler pathways, like EFA, though BANN pathways are more numerous, less polygenic, and require prior biological annotations [65].

There are several important limitations to our study. First, like linear regression, EFA can fit dozens to thousands of genetic variants (**Supplementary Figure S3**) but cannot jointly fit the entire genome. In practice, we pre-screen SNPs based on additive signals, such as selecting only GWAS-significant SNPs in UK Biobank, which enriches for SNPs with epistasis effects [24, 66]. Second, our polygenic epistasis tests do not establish interactions between any given SNP pair. In the future, sparse models (Bayesian or frequentist) could test which SNPs affect which EFs. Third, following [35], we have only studied four relatively simple complex human traits, and it will be important in the future to evaluate EFA more broadly across more complex traits such as height and BMI.

Another set of concerns are statistical false positives from phenotype scale and/or complex LD patterns. First, it is obvious that scale transformations of an additive trait can induce interactions [33, 45, 67, 68].

Nonetheless, 2/4 of our human EFA signals are qualitatively identical on the natural and quantile-normalized scales, and coordination is theoretically robust to modest rescaling [33]. Second, LD with unmeasured causal variants can cause dramatic and replicating false positives, especially in small *cis*-windows with large effects [18, 48–51]. We partly address this by LD-pruning SNPs, and we also use simulations to show that EFA’s polygenic nature reduces sensitivity to locus-specific LD biases. More broadly, EFA’s utility for prediction and biological characterization persists even if EFA signals reflect scaling and/or subtle LD patterns. Overall, these caveats are crucial for interpreting interactions, but they do not impact our central conclusions.

There are several extensions to EFA that may prove useful. First, EFA could jointly model multiple traits that partly share EFs. This would add power to detect shared EFs and provide a rich decomposition of pleiotropy that goes beyond genetic correlation, which is overly simplistic [69–73]. Second, EFA could model complex diseases in a generalized linear model framework, which is closely related to the idea of limiting disease pathways [34] and disease subtyping [57]. However, binary disease traits will have lower power and also have subtle scale dependence issues [74]. Third, we have focused on SNP interactions, but in principle, EFA is capable of modeling more powerful and/or interpretable genetic features such as imputed gene expression [75, 76], copy number variants [27, 28, 77], or polygenic scores for secondary traits [78] or exposures [41]. EFA could also incorporate non-genetic variables such as epigenomic marks, medical image-derived features [79], disease symptoms, or comorbidities from electronic health records. Ultimately, we hope that the EFA model and its strong results in real data provide a solid step toward unravelling epistasis in complex human traits.

## Code availability

A python implementation of EFA is available at: https://github.com/tangdavid/efa. This link also contains the code for all analyses in this paper, though the UKB analysis is not fully reproducible because the raw data is not public. An R implementation is available at: https://github.com/andywdahl/EFA.

## Data availability

We downloaded the first yeast data set from http://genomics-pubs.princeton.edu/YeastCross_BYxRM/data/cross.Rdata and the second yeast dataset from https://static-content.springer.com/esm/art%3A10.1038%2Fncomms9712/MediaObjects/41467_2015_BFncomms9712_MOESM729_ESM.zip

This research has been conducted using the UK Biobank Resource under application number 30397.

## Author contributions

D.T. developed statistical methodology, performed analysis, and wrote the manuscript. J.F. performed analysis. A.D. conceived and supervised the project and wrote the manuscript.

## Acknowledgements

We thank Matthew Stephens, Sasha Gusev, Sriram Sankararaman, and Noah Zaitlen for helpful feedback. We also thank the participants in UKB for making this study possible. Finally, we thank the Center for Research Informatics and the Research Computing Center for providing the compute resources necessary for carrying out this project. A.D. is supported by K25HL157603.

## Methods

### Mathematical description of Epistasis Factor Analysis

Epistasis Factor Analysis (EFA) aims to find a low-dimensional representation of polygenic epistasis that is both statistically powerful and biologically interpretable. The EFA model is:

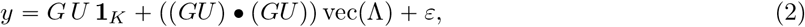

where *y* is a quantitative trait measured on *N* samples, *G* is an *N* × *M* matrix of genotypes at *M* SNPs, *U* is an *M* × *K* matrix of SNP effects on each of *K* factors, Λ is a symmetric *K* × *K* matrix of factor-level interactions, • is the face-splitting product (*A* •*B* is a matrix where each column is an element-wise product of a column of *A* and a column of *B*), and *ε* is independent and identically distributed (i.i.d.) Gaussian noise.

Intuitively, each column of *U* represents the SNP weights on a different latent epistasis factor (or EF), and *P*_,*k*_ := *GU*_,*k*_ is an individual’s weight on the *k*-th EF. Λ_*kk′*_ represents the factor-level interaction between EFs *k* and *k*. When Λ_*kk*_′ > 0, factors *k* and *k* ′ interact synergistically, meaning that the their combined phenotypic effect is greater than the sum of their individual effects. The reverse is true for antagonistic factors, when Λ_*kk*_′ > 0. When Λ_*kk*_′ = 0, then factors *k* and *k* do not interact. Finally, Λ_*kk*_ refers to the quadratic effect of pathway *k*, which can help identify the pathway weights in *U* and which we do consider *bona fide* statistical epistasis. Nonetheless, our primary interest is in the Λ_*kk*_′ terms where *k* ≠ *k*, as we are motivated by unravelling interactions across distinct biological factors.

In the main text, we call this general model EFA^*^ and focus instead on the special case where (i) *K* = 2 and (ii) Λ_11_ = Λ_22_ = 0. We focus on this special case for simplicity and also because it is identified, i.e., the estimated EFs and their interactions are meaningful (**Supplement Section 1.5.1**). In contrast, the general EFA^*^ model in (2) is not identified without further assumptions, such as that *U* is sparse (Note that most factor models, including PCA, are not identified without additional assumptions.) We prove these facts and comprehensively characterize the theoretical properties of EFA in the **Supplement**. We also formally characterize the equivalence class of solutions to EFA*, proving that the sign of Λ_*kk*_′ is well-identified and providing a canonical representative of the equivalence class for *K* = 2 (**Supplement Section 1.5.2**). We also calculate the level of coordinated epistasis under EFA in the **Supplement** [33].

EFA is a special case of the standard pairwise polygenic epistasis model that underlies many epistasis models for complex traits:

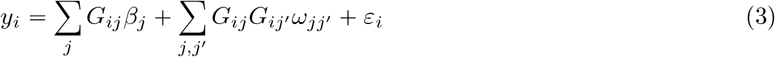

where *β*_*j*_ is the additive effect of SNP *j*, and *ω*_*jj*_′ is the epistatic interaction between SNPs *j* and *j*. The factor-level interaction model in EFA can be directly connected with this SNP-level epistasis model by the linked factorization of the SNP-level additive and epistasis effects:

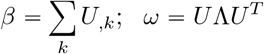

### Fitting EFA parameters

We always estimate the parameters of EFA using maximum likelihood (**Supplement Section 1.4**). In general, we fit the EFA^*^ model using gradient descent based on automatic differentiation. We evaluate multiple random restarts, each initialized with *U*_,*k*_ drawn from a neighborhood around 0. We use this approach in simulations and in the yeast analyses. This generic, flexible estimation method can easily accommodate future model extensions, e.g., introducing sparsity in *U* with *ℓ*_1_ penalties or “anchoring” EFs to specific SNPs (**Supplement Section 1.5.3**).

To scale our method to UKB sample size, we derive a block coordinate ascent algorithm (**Supplement Section 1.4**). The updates are remarkably simple, requiring only iterative regressions that scale linearly in sample size and quadratically in SNPs (i.e., *O*(*NM* ^2^)). The key to this efficiency is that we only evaluate EF-EF interactions and never explicitly evaluate SNP-SNP interactions, which would cost *O*(*NM* ^4^). With 100 SNPs, as in our UKB analyses, this is a 10,000-fold speedup.

### Uncoordinated models of polygenic epistasis

We compare EFA with other polygenic models in simulations and real data analyses. All comparison methods are special cases of the overarching polygenic pairwise epistasis model in (3).

First, we compare to the standard additive model, which ignores epistasis by setting *ω*_*jj*_′ = 0 for all *j, j*. Because we mostly study scenarios with *M* < *N*, we primarily fit the additive model with fixed effects (by ordinary least squares). The predictions from this model are then essentially Polygenic Risk Scores (PRS). We also fit the additive model with random effects (assuming each *β*_*j*_ is i.i.d. Gaussian) in the real yeast analysis, enabling us to fit the full genome-wide data without any pruning.

Second, we compare to the standard pairwise epistasis model in (3) fit with fixed-effects. Because this model has *O*(*M* ^2^) parameters, jointly fitting all pairwise epistasis effects is often noisy or even impossible. Instead, we fit each pairwise interaction *ω*_*jj*_′ one-at-a-time (for *j* ⩽ *j*) while controlling for additive fixed effects (note that only *ω*_*jj*_′ ⩽ *ω*_*j*_′ _*j*_ is identified).

Third, we compare to the uncoordinated random effect model for pairwise epistasis [31]. This model assumes that *ω*_*jj*_′ are drawn i.i.d. Gaussian. This assumption simplifies computation because the *ω* can be easily marginalized out of the likelihood. We fit the variance components (one for *β* and one for *ω*) with maximum likelihood. We derive the best linear unbiased predictors (BLUPs) for both *ω* and for phenotype prediction in the **Supplement**.

Finally, we also develop a novel approach to estimate latent pathways from uncoordinated estimates by applying PCA to the estimated *ω* matrix. This is a naive baseline approach that only involves post-processing uncoordinated estiamtes. In contrast, EFA directly learns pathways, linking the low dimensional components of *ω* with the additive effects.

### Simulations

We use a polygenic simulation framework that is fully described in the **Supplement Section 3**. In brief, we simulate under the EFA model (2) with independent SNPs. The latent pathways, *U*_,*k*_, are drawn from i.i.d. Gaussians with variance chosen to give the appropriate additive heritability (recall that the additive effect is related to the latent pathways by *β* = Σ _*k*_ *U*_,*k*_). The upper triangular entries of Λ are also drawn fro i.i.d. Gaussians with variance chosen to give the appropriate epistasis heritability. The pairwise polygenic epistasis effects are then generated by 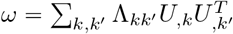.

We also simulate partly coordinated epistasis by adding i.i.d. Gaussian random variables to each entry of a coordinated *ω*. By varying the relative contribution of these effects, we can interpolate between the EFA model (coordinated) and the classical random effect model (uncoordinated). By choosing the relative variances of the different components appropriately, we are able to simulate the standard additive model, the standard uncoordinated model, and the coordinated EFA model.

Finally, we use two alternative simulation frameworks for evaluating the calibration of our bootstrap test with respect to LD and dominance that we describe fully in the **Supplement**.

### Yeast data

We downloaded genotype and phenotype data for prototrophic haploid segregants from a cross between a laboratory strain and a wine strain of yeast [13]. This dataset consisted of 46 quantitative traits for 1,008 samples genotyped at 11,623 unique markers. We standardized each phenotype and SNP to mean 0 and variance 1. To account for linkage disequilibrium in fixed effect models, we clumped+thresholded variants using threshold of *r*^2^ < 0.2, minimum distance of 250 kb, and marginal additive *p* < 0.05 using PLINK 1.9 [80]. We performed clumping+thresholding separately on each cross-validated fold to avoid bias from overfitting [81].

Additionally, we downloaded a second yeast growth trait dataset from the same group with a larger sample size (*N* = 4390) [14]. This second dataset contained 20/46 of the growth phenotypes in the first dataset, and it included 28,220 unique genotype markers for each sample. We preprocessed this dataset in the same way as above.

We evaluated phenotype prediction accuracy with 10-fold cross validation, using the same 10 folds across all methods so we could test for differential prediction accuracy with paired *t*-tests.

### UK Biobank data

We studied the same four traits in UK Biobank (UKB) that were extensively characterized in [35]: urate, IFG1, testosterone in males, and testosterone in females. We analyzed unrelated “white British” individuals as defined by UKB. We used the top 100 LD-pruned GWAS hits from [35], or all 77 GWAS hits for female testosterone. Prior to fitting EFA, we regressed out sex, age, batch, assessment center, and the top 10 genotype PCs from the phenotype.

We assess statistical significance using 499 bootstrap samples to estimate standard errors and confidence intervals on the pathway level interaction. We compute empirical *p*-values as 2 · min(*B*_*λ*<0_, *B*_*λ*>0_) where *B*_*λ*<0_ is the number of bootstrap samples with *λ* < 0 and *B*_*λ*>0_ is the number of bootstrap samples with *λ* < 0. We also report Gaussian *p*-values obtained from *z*-scores computed as the bootstrap mean divided by the bootstrap standard deviation.

To test biological significance, we used the annotations of GWAS hits to biological pathways provided in [35]. We assigned SNPs to the epistasis factors by (1) polarizing SNPs to have positive additive effect sizes and (2) choosing the largest 15 effects on each pathway after subtracting out additive effects from the pathways. To test robustness, we also evaluate results using 15 or 20 SNPs per pathway (**Supplementary Table 3-7**). We calculate *p*-values for enrichment of biological pathways in epistasis factors with a hypergeometric test.

