## Supplement for "Factorizing polygenic epistasis improves prediction and uncovers biological pathways in complex traits"

**4 Supplementary Figures** **9**

**5 Supplementary Tables** **17**

### 1 Analytical Properties of EFA

#### 1.1 Generalizing EFA to more pathways

Our main text focuses on the case of two EFs. Here, we provide the more general EFA model for an arbitrary number of pathways, $K$. In the main text, we call this model EFA* for clarity, but we just call it EFA in this Supplement. The general EFA model can be formally written as:

$$y_i = \sum_k P_{ik} + \sum_{k,k'} \Lambda_{kk'} P_{ik} P_{ik'} + \varepsilon_i; \qquad P_{ik} := \sum_j G_{ij} U_{jk} \tag{1}$$

where $y$ is the quantitative complex trait, $G$ is the genotype matrix, $P$ is the matrix of pathways, $U$ is the matrix of loadings of each SNP onto each pathway, and $\varepsilon$ is assumed to be i.i.d. Gaussian noise. Note that this model also includes self-interaction terms, i.e., $\Lambda_{kk}$. In the main text, we assume $K = 2$ and also that $\Lambda_{11} = \Lambda_{22} = 0$. This reduces the interaction terms down to just $\Lambda_{12}$, which we call $\lambda$ in the main text for simplicity.

We loosely refer to both $P$ and $U$ as Epistasis Factors (EFs), akin to how principal components can refer to both the left and right singular vectors of a matrix.

Note that $\omega$ is only identified modulo symmetrization, i.e., $\omega$ and $\omega^T$ are equivalent. If desired, we can break this equivalence class by requiring $\omega$ to be symmetric, or to be (non-strictly) lower-triangular. In the case of the simplified EFA model, with $K = 2$ pathways and no self-interaction, $\omega$ is identified up to transpoition, which is equivalent to switching the labels of EF1 and EF2.

In principle, it is straightforward to modify this objective function to accommodate heteroscedastic and/or non-Gaussian noise terms, $\epsilon$. In particular, this easily generalizes to generalized linear models, such as logistic regression, which can enable EFA to fit binary disease traits. Nonetheless, while these generalizations are straightforward computationally, they remain challenging from a statistical perspective, especially in terms of appropriately choosing link functions for binary traits [2].

#### 1.2 Connecting EFA with genome-wide pairwise epistasis

In the main text, we define the standard genome-wide pairwise epistasis model as:

$$y_i = \sum_s G_{is}\beta_s + \sum_{s,s'} \omega_{ss'} G_{is} G_{is'} + \varepsilon_i \tag{2}$$

Conceptually, this closely resembles the EFA model except the interactions occur at the level of SNPs ($\omega_{ss'}$) rather than the level of pathways ($\Lambda_{kk'}$).

This model is identical to the EFA model if we assume a specific additional structure on $\omega$:

$$\beta = \sum_k U_{,k}; \quad \omega = U \Lambda U^T \tag{3}$$

That is, these models coincide if genome-wide epistasis solely reflects interactions between $K$ additive pathways. Note that non-interacting additive effects can be included without loss of generality using a 'sink' pathway, i.e., defining the $K$-th pathway as the remaining additive effects not captured by interacting pathways: $U_{,K} := \beta - \sum_{k<K} U_{,k}$. Thus, the key assumption is not that all SNPs contribute to interacting pathways; rather, the assumption is that all epistasis is explained by $K$ pathway-level interactions.

There are two key differences between EFA and our *post hoc* approach that applies PCA to Uncoordinated $\omega$ estimates. First, EFA explicitly learns a low-dimensional representation to $\omega$, which is more efficient than a two-step approach. This is the reason that EFA performs better in estimating $\omega$ even when additive effects are absent (main Figure 2a). However, the more important difference is that EFA links together additive effects with epistasis effects, which can be seen by the fact that EFA uses the same $U$ term to jointly partition the additive effects $\beta$ and the epistasis effects $\omega$ in (3).

#### 1.3 Connecting EFA to Coordinated Epistasis

EFA is essentially an explicit model of the pathway-level epistasis model that motivated Coordinated Epistasis in [3]. To flesh out this connection, we derive here the exact level of Coordination ($\gamma$) as defined in [3] under the EFA model. That is, we calculate:

$$\gamma := \mathrm{Cov}_{j \neq j'}(\beta_j \beta_{j'}, \Omega_{jj'}) = \mathrm{Cov}_{j \neq j'} \left( \sum_{k=1}^{K} U_{jk} \cdot \sum_{k'=1}^{K} U_{j'k'}, \sum_{k,k'=1}^{K} \Lambda_{kk'} U_{jk} U_{j'k'} \right)$$

The first equality is the definition of CE from [3]; the second uses EFA's assumptions on $\beta$ and $\omega$ in (3).

For simplicity, we assume that pathway weights $U_{jk}$ are random and independent, with $\mathbb{E}(U_{jk}) = 0$ and

$\mathbb{V}(U_{jk}) = \sigma_k^2$. Then:

$$\begin{aligned}\gamma &= \sum_{k,k',p,p'} \Lambda_{kk'} \text{Cov}_{j \neq j'} \left(U_{jp} U_{j'p'}, U_{jk} U_{j'k'}\right) \\ &= \sum_{k,k',p,p'} \Lambda_{kk'} \mathbb{E}_{j \neq j'} \left(U_{jp} U_{j'p'} U_{jk} U_{j'k'}\right) \\ &= \sum_{k,k'} \Lambda_{kk'} \mathbb{V}(U_{jk}) \mathbb{V}(U_{j'k'}) \\ &= \sum_{k,k'} \sigma_k^2 \sigma_{k'}^2 \Lambda_{kk'}\end{aligned} \tag{4}$$

Intuitively, this last equality says that the overall Coordination is an average of the pairwise pathway-level interaction terms ($\Lambda_{kk'}$) weighted by the pathway strengths ($\sigma_k^2$). (This calculation can easily be generalized to allow correlations between SNPs and/or between pathway weights for a single SNP, though the result is less intuitive.)

#### 1.4 Model Fitting

We fit the latent pathways in our model with maximum likelihood. Assuming normal error terms, the maximum likelihood estimate is equivalently formulated as the parameter estimate that minimizes the squared error loss, i.e.,

$$\hat{U}, \hat{\Lambda} = \arg\min_{U,\Lambda} L(Y, G; U, \Lambda) = \arg\min_{U,\Lambda} \|Y - (GU\mathbf{1}_K + (G \bullet G)(U \otimes U)\text{vec}(\Lambda))\|_2^2 \tag{5}$$

where $1_K$ is a vector of 1s and $\bullet$ is the face-splitting product. Concretely, $G \bullet G$ is the matrix containing all products of all pairs of columns in $G$. This seems somewhat complicated, but it is just the matrix of interactions in a linear model; fitting interaction terms in a linear model always implicitly creates such a matrix.

We employ two optimization strategies to compute the MLEs. Our default approach is to use generic gradient-based optimization techniques. Specifically, we use the Adam optimizer implemented in PyTorch's `torch.optim.Adam` class. We run the optimization procedure with random restarts and then choose the result with the smallest squared error. By default, we use 20 random restarts and initialize all parameters from a centered normal distribution with variance 0.1.

Our second approach iterates analytic block coordinate ascent steps but only applies to the special case with no self-interactions, i.e., $\Lambda_{kk} = 0$ for all $k$. In this case, we sequentially find the conditional maximum likelihood estimates for each block of parameter coordinates and iterate. Algorithmically, we iterate the following sequence of updates until convergence. We first update the pathways as

$$\begin{aligned}U_1^{(t+1)} &\leftarrow \arg\min_{U_1} L(Y, G; U_1, U_2^{(t)}, \ldots, U_K^{(t)}, \Lambda^{(t)}) \\ U_2^{(t+1)} &\leftarrow \arg\min_{U_2} L(Y, G; U_1^{(t+1)}, U_2, \ldots, U_K^{(t)}, \Lambda^{(t)}) \\ &\vdots \\ U_K^{(t+1)} &\leftarrow \arg\min_{U_K} L(Y, G; U_1^{(t+1)}, U_2^{(t+1)}, \ldots, U_K, \Lambda^{(t)})\end{aligned}$$

We then sequentially update the interaction weights as

$$\begin{aligned}\Lambda_{12}^{(t+1)} &\leftarrow \arg\min_{\Lambda_{12}} L(Y, G; U^{(t+1)}, \Lambda_{12}, \Lambda_{13}^{(t)}, \ldots, \Lambda_{(K-1)K}^{(t)}) \\ \Lambda_{13}^{(t+1)} &\leftarrow \arg\min_{\Lambda_{13}} L(Y, G; U^{(t+1)}, \Lambda_{12}^{(t+1)}, \Lambda_{13}, \ldots, \Lambda_{(K-1)K}^{(t)}) \\ &\vdots \\ \Lambda_{(K-1)K}^{(t+1)} &\leftarrow \arg\min_{\Lambda_{(K-1)K}} L(Y, G; U^{(t+1)}, \Lambda_{12}^{(t+1)}, \Lambda_{13}^{(t+1)}, \ldots, \Lambda_{(K-1)K})\end{aligned}$$

Given all other parameters, we can rewrite the objective as a simple quadratic function of $U_k$:

$$L = \|Y - \left(\sum_i GU_i + \sum_{i,j} \Lambda_{ij}(GU_i) \circ (GU_j)\right)\|_2^2 = \|(Y - C_k) - A_k U_k\|_2^2$$

with

$$A_k = \left(I + 2\text{diag}\left(\sum_{i \neq k} \Lambda_{ik} GU_i\right)\right) G \quad \text{and} \quad C_k = \sum_{i,j \neq k} \Lambda_{ij} GU_i \circ GU_j + \sum_{i \neq k} GU_i \tag{6}$$

This can immediately be solved by ordinary least squares:

$$\arg\min_{U_k} L = (A_k^T A_k)^{-1} A_k^T (Y - C_k) \tag{7}$$

Likewise, to update $\Lambda_{pq}$, we can rewrite the objective as

$$L = \|(Y - C_{pq}) - 2\Lambda_{pq} X_{pq}\|^2$$

where the vector $X_{pq}$ and the constant $C_{pq}$ do not depend on $\Lambda_{pq}$:

$$X_{pq} = (GU_p) \circ (GU_q) \quad \text{and} \quad C_{pq} = \sum_{i,j:(i,j) \neq (p,q)} \Lambda_{ij}(GU_i) \circ (GU_j) + \sum_i GU_i \tag{8}$$

The least squares solution is then:

$$\arg\min_{\Lambda_{pq}} L = \frac{X_{pq}^T (Y - C_{pq})}{\|X_{pq}\|_2^2} \tag{9}$$

Note that the updates for $U_k$ and $\Lambda_{pq}$ appear to require multiplying $N \times N$ matrices with $N \times 1$ vectors, these matrices are diagonal and hence all operations can be performed in $O(NS^2)$ time rather than $O(N^2S^2)$ time. This is essential for scaling EFA to large datasets.

In practice, we find that the coordinate descent algorithm behaves well when we initialize near the additive model, i.e., $\Lambda = 0$ and $U_k = \hat{\beta}/K$. We iterate the algorithm until convergence; if it does not converge within 5,000 iterations, we restart after adding small random noise around the initialization.

#### 1.5 Identification

The general EFA model is not identified, i.e., the solution to the maximum likelihood problem in (5) is not unique. Specifically, the likelihood is invariant to the following transformation:

$$(U, \Lambda) \mapsto (UA^{-1}, A\Lambda A^T)$$

for any invertible matrix $A \in \mathbb{R}^{K \times K}$ with rows that sum to 1. In other words, if $(U, \Lambda)$ is a maximum likelihood solution, then so is $(V, \Lambda') := (UA^{-1}, A\Lambda A^T)$. This is more severe than the standard label-switcghing identification in factor models because the solution is not a discrete set of functionally-identical answers but, rather, a continuous manifold of solutions with no clear interpretation.

To see this equivalence, first observe that:

$$GV\mathbf{1}_K = GUA^{-1}\mathbf{1}_K = GU\mathbf{1}_K \tag{10}$$

This shows that our transformation preserves $\beta$. Second, observe that:

$$\begin{aligned}
(G \bullet G)(V \otimes V)\text{vec}(\Lambda') &= (G \bullet G)(UA^{-1} \otimes UA^{-1})\text{vec}(A\Lambda A^T) \\
&= (G \bullet G)(U \otimes U)(A^{-1} \otimes A^{-1})\text{vec}(A\Lambda A^T) \\
&= (G \bullet G)(U \otimes U)\text{vec}(A^{-1}A\Lambda A^T(A^T)^{-1}) \\
&= (G \bullet G)(U \otimes U)\text{vec}(\Lambda)
\end{aligned} \tag{11}$$

This shows our transformation preserves $\omega$.

Because the likelihood depends only on $\beta$ and $\omega$, and because our transformation by $A$ preserves these terms, our transformation does not change the likelihood. Thus, the span of this transformation (by an invertible matrix $A$ with row sums of 1) defines a class of equivalent solutions to EFA.

This is problematic because it means the general EFA model does not provide meaningful EFs, because there are infinitely many sets of EFs that explain the data equally well. Nonetheless, the estimates of $\omega$ and the predictions are identified; this is why we evaluate EFA* only in these terms in the main text.

However, there are special cases of the general EFA model where we can achieve meaningful identification.

##### 1.5.1 Special Case 1: K=2, No Self-Interactions

When $K = 2$ and we restrict the diagonal entries of $\Lambda$ to zero, the optimization problem is well-identified (modulo label switching). Mathematically, this follows from the fact that the only matrix $A$ in our equivalence class above is the identity.

##### 1.5.2 Special Case 2: Canonical Representatives with Self-Interaction

Depending on conditions involving the overall sign of $\gamma$, we can uniquely choose canonical representatives of the equivalence class that only have self interactions between latent pathways (models with diagonal $\Lambda$). For the sake of simplicity, we restrict ourselves to the case where $K = 2$ (similar ideas extend inductively with considerable casework to $K > 2$).

Specifically, we desire to show that for any $\Lambda$, there exists a unique (up to permutation) and equivalent $\Lambda'$ with $\Lambda'_{12} = 0$ that satisfies certain conditions for $\Lambda'_{11}$ and $\Lambda'_{22}$. Without loss of generality, we may assume that $\Lambda$ is diagonal. This is because for any $\Lambda = (\lambda_{ij})$, we can obtain an equivalent diagonal $\Lambda' = A\Lambda A^T$ where

$$A = \begin{pmatrix} 1 & 0 \\ \frac{\lambda_{21}}{\lambda_{21}-\lambda_{11}} & 1 - \frac{\lambda_{21}}{\lambda_{21}-\lambda_{11}} \end{pmatrix}$$

For any diagonal $\Lambda$ and invertible 2x2 matrix $A$ with $A1 = 1$, we have

$$\Lambda = \begin{pmatrix} \lambda_1 & 0 \\ 0 & \lambda_2 \end{pmatrix} \text{ and } A = \begin{pmatrix} a & 1-a \\ 1-b & b \end{pmatrix}$$
$$\implies \Lambda' = A\Lambda A^T = \begin{pmatrix} a^2\lambda_1 + (1-a)^2\lambda_2 & a(1-b)\lambda_1 + b(1-a)\lambda_2 \\ \cdot & b^2\lambda_2 + (1-b)^2\lambda_1 \end{pmatrix}$$

For $\Lambda'$ to be diagonal, we require

$$a = \frac{-\lambda_2 b}{\lambda_1(1-b) - \lambda_2 b} \iff A = \begin{pmatrix} \frac{-\lambda_2 b}{\lambda_1(1-b)-\lambda_2 b} & \frac{\lambda_1(1-b)}{\lambda_1(1-b)-\lambda_2 b} \\ 1-b & b \end{pmatrix}$$

In turn, we can write $\Lambda'$ as follows

$$\begin{aligned} \Lambda' := A\Lambda A^T &= \begin{pmatrix} \left(\frac{1}{\lambda_1(1-b)-\lambda_2 b}\right)^2 \left(b^2\lambda_2^2\lambda_1 + \lambda_1^2(1-b)^2\lambda_2\right) & 0 \\ 0 & b^2\lambda_2 + (1-b)^2\lambda_1 \end{pmatrix} \\ &= \begin{pmatrix} \frac{\lambda_1\lambda_2}{(\lambda_1(1-b)-\lambda_2 b)^2} \left(b^2\lambda_2 + (1-b)^2\lambda_1\right) & 0 \\ 0 & b^2\lambda_2 + (1-b)^2\lambda_1 \end{pmatrix} \\ &= \left(b^2\lambda_2 + (1-b)^2\lambda_1\right) \begin{pmatrix} \frac{\lambda_1\lambda_2}{(\lambda_1(1-b)-\lambda_2 b)^2} & 0 \\ 0 & 1 \end{pmatrix} \end{aligned}$$

As it will soon become clear, choosing a canonical representative splits naturally into two cases: $\lambda_1\lambda_2 > 0$ and $\lambda_1\lambda_2 < 0$.

In the first case with $\lambda_1\lambda_2 > 0$, we can choose a matrix $A$ such that $\Lambda'_{11} = \Lambda'_{22}$. This is equivalent to choosing some scalar $b$ such that

$$\frac{\lambda_1\lambda_2}{(\lambda_1(1-b) - \lambda_2 b)^2} = 1 \iff \lambda_1\lambda_2 = (\lambda_1(1-b) - \lambda_2 b)^2$$

Solving for $b$ gives the following solution

$$b = \frac{\lambda_1 \pm \sqrt{\lambda_1\lambda_2}}{\lambda_1 + \lambda_2} \tag{12}$$

It is now clear why the problem of choosing a canonical representative naturally splits into two cases. When $\lambda_1\lambda_2 > 0$, we can always uniquely project (up to label switching) $\Lambda$ into a diagonal $\Lambda'$ that is a scalar multiple of the identity matrix. However, it is never possible to project $\Lambda$ into such a canonical representative when $\lambda_1\lambda_2 < 0$ .

In the second case, we can instead project $\Lambda$ into a diagonal matrix $\Lambda'$ with $\Lambda'_{22} = \pm 1$, where the sign depends on whether or not $\lambda_1 + \lambda_2 > \lambda_1\lambda_2$ or $\lambda_1 + \lambda_2 < \lambda_1\lambda_2$. Specifically, we are choosing some scalar $b$ such that

$$b^2\lambda_2 + (1-b)^2\lambda_1 = c$$

where $c$ is either 1 or $-1$. Solving, we get that

$$b = \frac{\lambda_1 \pm \sqrt{c\lambda_1 + c\lambda_2 - \lambda_1\lambda_2}}{\lambda_1 + \lambda_2} \tag{13}$$

This shows that we can always project $\Lambda$ into an equivalent $\Lambda'$ such that $\Lambda'_{22}$ is either 1 or $-1$ depending on whether $\lambda_1 + \lambda_2 > 0$ or $\lambda_1 + \lambda_2 < 0$ respectively.

While these canonical representatives are somewhat removed from the motivating biology, this exercise demonstrates that the signed direction of coordinated interactions is an intrinsic property that affects the qualitative nature of the model. Specifically, there is a sense in which systems with interactions that have opposite effects are fundamentally different from systems where all interactions have the same directional effect; the first set of models projects to one class of canonical representatives (case 2) while the second set projects to another class of canonical representatives (case 1).

Moreover, when the the signs of the interactions are opposed, the canonical representatives again split into two categories depending on whether the overall sign of the interactions, $\lambda_1 + \lambda_2$, is positive or negative. This demonstrates the mathematical value in understanding the directional effects of coordinated interactions.

##### 1.5.3 Special Case 3: Anchor SNPs

In the general case of the model, we can leverage anchor SNPs to get identified estimates. Anchor SNPs are SNPs that we can reasonably expect to only affect one pathway, providing a biological prior to help break the equivalence relation.

Mathematically, if we have $K$ different pathways such that $U \in \mathbb{R}^K$, we can define a set of $K$ SNPs indexed as $\{a_1, \ldots, a_K\}$ such that

$$U_{a_j,p} = \begin{cases} x_j \neq 0 & j = p \\ 0 & \text{otherwise} \end{cases} \tag{14}$$

We claim that this constraint breaks the equivalence relation. Suppose we have two equivalent solutions $(V, \Lambda')$ and $(U, \Lambda)$ satisfying this constraint. As characterized above, $V$ and $U$ are related by $VA = U$ for some invertible matrix $A$. For any $j \in \{a_1, \ldots, a_k\}$, we have that

$$\sum_{p=1}^{k} V_{j,p} A_{p,j} = x_j = U_{j,p} \text{ and } \sum_{p=1}^{k} V_{j,p} A_{p,l} = 0 = U_{l,p} \text{ for } l \neq j \tag{15}$$

Using the constraints on $V$, this simplifies to

$$V_{j,j} A_{j,j} = x_j \text{ and } V_{j,j} A_{j,l} = 0 \text{ for } l \neq j \tag{16}$$

This is only possible if $A$ is the identity matrix, showing that the presence of anchor SNPs allows us to resolve the identifiability of the model.

### 2 Random Effects BLUPs

We now derive the random effect best linear unbiased predictors (BLUPs) of $\beta$ and $\omega$ for the uncoordinated epistasis model. We assume that $\beta$ and $\omega$ are drawn from independent normal distributions, i.e.,

$$y = G\beta + (G \bullet G)\omega + \varepsilon$$
$$\beta \sim N(0, \frac{\sigma_\beta^2}{M} I) \text{ and } \omega \sim N(0, \frac{\sigma_\omega^2}{M^2} I)$$

Then the vector $(\beta, \omega, y)$ follows a multivariate normal distribution with the following covariance.

$$(\beta, \omega, y) \sim N\left(0, \begin{pmatrix} \frac{\sigma_\beta^2}{M} I & 0 & \frac{\sigma_\beta^2}{M} G^T \\ 0 & \frac{\sigma_\omega^2}{M^2} I & \frac{\sigma_\omega^2}{M^2} (G \bullet G)^T \\ \frac{\sigma_\beta^2}{M} G & \frac{\sigma_\omega^2}{M^2} (G \bullet G) & \sigma_\beta^2 K + \sigma_\omega^2 K \circ K + \sigma_\varepsilon^2 I \end{pmatrix}\right)$$

The conditional distribution of multivariate normals then gives us that

$$\mathbb{E}[\beta | y] = \left(\frac{\sigma_\beta^2}{M} G\right)^T \left(\sigma_\beta^2 K + \sigma_\omega^2 K \circ K + \sigma_\epsilon^2 I\right)^{-1} y \tag{17}$$

and

$$\mathbb{E}[\omega | y] = \left(\frac{\sigma_\omega^2}{M^2} G \bullet G\right)^T \left(\sigma_\beta^2 K + \sigma_\omega^2 K \circ K + \sigma_\epsilon^2 I\right)^{-1} y \tag{18}$$

### 3 Simulation Details

#### 3.1 Base Simulations

We start by simulating the genotypes of independent individuals. For each locus $j$, we draw the allele frequency $f_j$ i.i.d from a Beta(2, 2) distribution. We then independently draw the genotype for each individual $i$ at locus $j$ by $G_{ij} \sim Binom(2, f_j)$. We standardize the genotypes at each locus to have mean zero and variance 1. We adjust the epistatic and additive heritability by scaling $U$ and $\Lambda$ appropriately. For the sake of simplicity and interpretability, we scale all parameters such that the phenotypes have variance 1.

To simulate partial coordination, we define the epistasis effects at the SNP level and add together coordinated and uncoordinated effects. The coordinated component is proportional to the EFA contribution: $\omega_{coord} = U\Lambda U^T$. The uncoordinated component is proportional to a random matrix whose elements are drawn i.i.d. Gaussian, $\omega_{uncoord}$. We then simulate the phenotype from (2) using $\omega := a\omega_{coord} + b\omega_{uncoord}$, where $a$ and $b$ define the fraction of $\omega$ that is coordinated.

For the base simulation parameters, we assume a perfectly coordinated model where the epistatic effect, additive effect, and individual level noise explain 25, 25, and 50 percent of the phenotypic variance respectively. In every case, we simulate 1,000 individuals and 20 SNPs.

#### 3.2 LD Simulations

To assess EFA's robustness to tagging LD, we consider an alternative simulation where all observed loci are non-causal but in LD with an unobserved causal locus. Specifically, we draw 10 triplets of loci from a trivariate normal distribution:

$$(G_1, G_2, G_3)^T \sim N\left(0, \begin{pmatrix} 1 & r & r \\ r & 1 & 0 \\ r & 0 & 1 \end{pmatrix}\right) \tag{19}$$

We choose the maximum $r = 0.7$ such that the covariance matrix is positive semi definite. For each triple, we take $G_1$ and $G_2$ as our non-causal observed loci, and draw a causal effect size $\beta \sim N(0,1)$ for $G_3$, our unobserved locus. The effect sizes are then scaled to give the appropriate level of heritability.

#### 3.3 Dominance Simulations

We also simulate under a dominance model to assess whether or not our results are driven purely by dominance effects. We use the same baseline simulation framework, but set the percent of coordinated epistasis to zero and simulate diagonal uncoordinated effects, i.e., $\omega_{uncoord}$ is diagonal.

### 4 Supplementary Figures

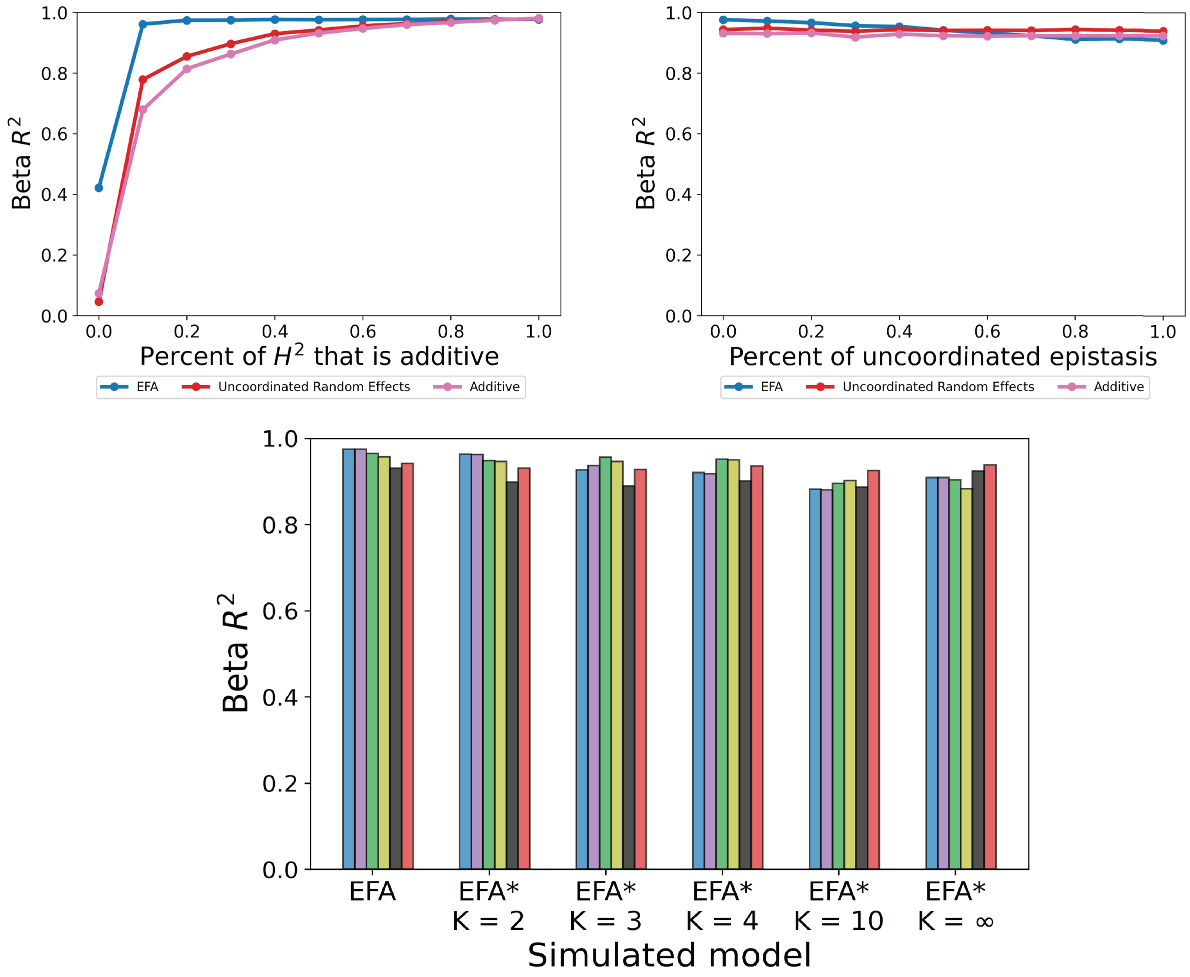

Figure S1: Estimation accuracy for additive effects. Simulations correspond to main text figure 2. EFA provides better estimates of additive effects than the additive model itself when there is sufficient epistasis heritability (top left), when it is mostly coordinated (top right), and when $K$ is chosen well (bottom).

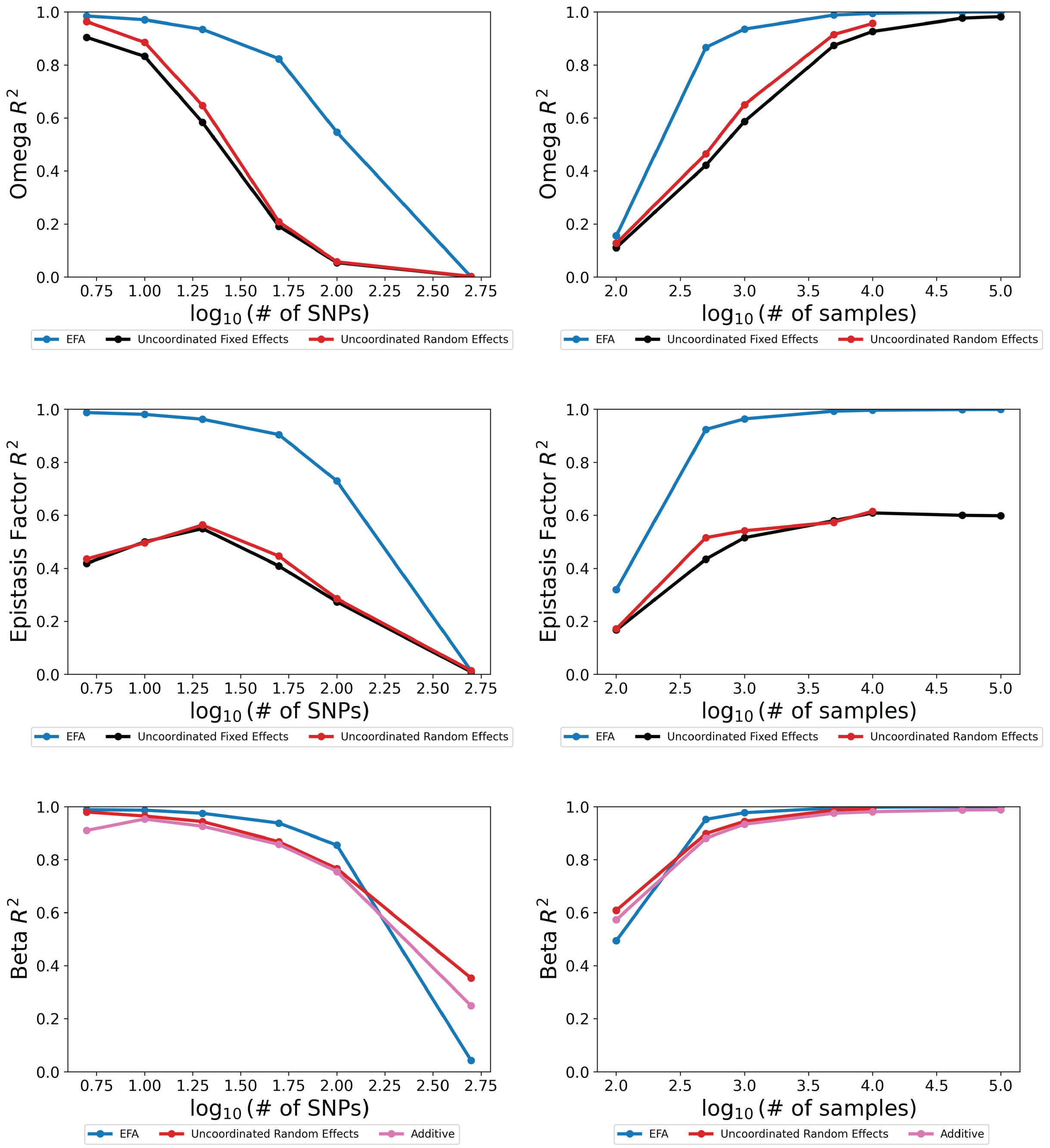

Figure S2: Estimation accuracy (defined as $R^2$ between the estimated and simulated parameters) in simulations varying sample size or number of SNPs. All methods improve with sample size and decrease with number of SNPs. Simulations varying the number of SNPs use a fixed sample size of $N = 1000$ and simulations varying sample size use a fixed number of SNPs of $M = 20$.

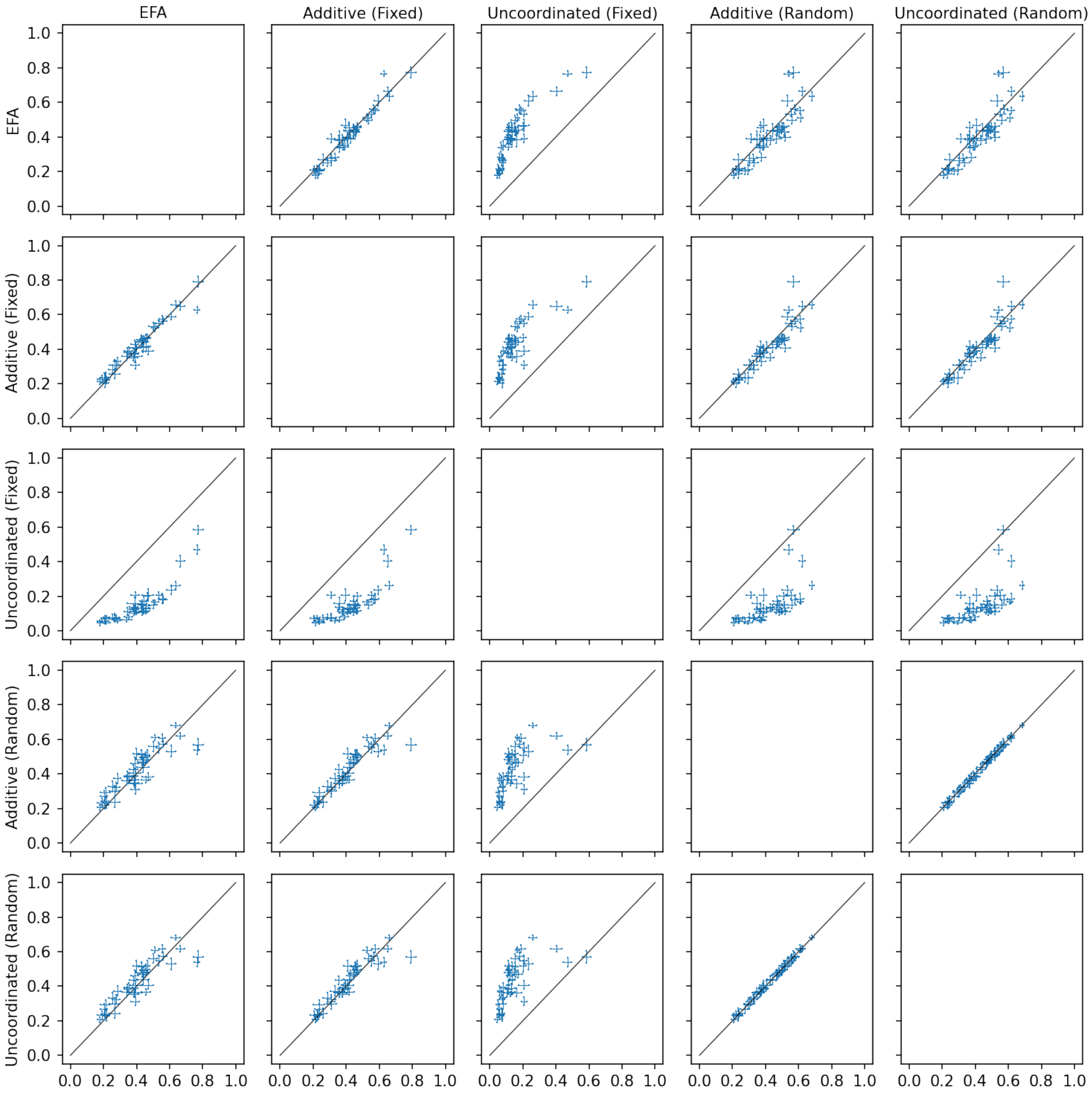

Figure S3: Pairwise comparisons of different polygenic prediction models for the 46 yeast traits in the first dataset. Prediction is measured by the average $R^2$ between the out-of-sample predicted phenotypes and the true phenotypes in 10-fold cross validation.

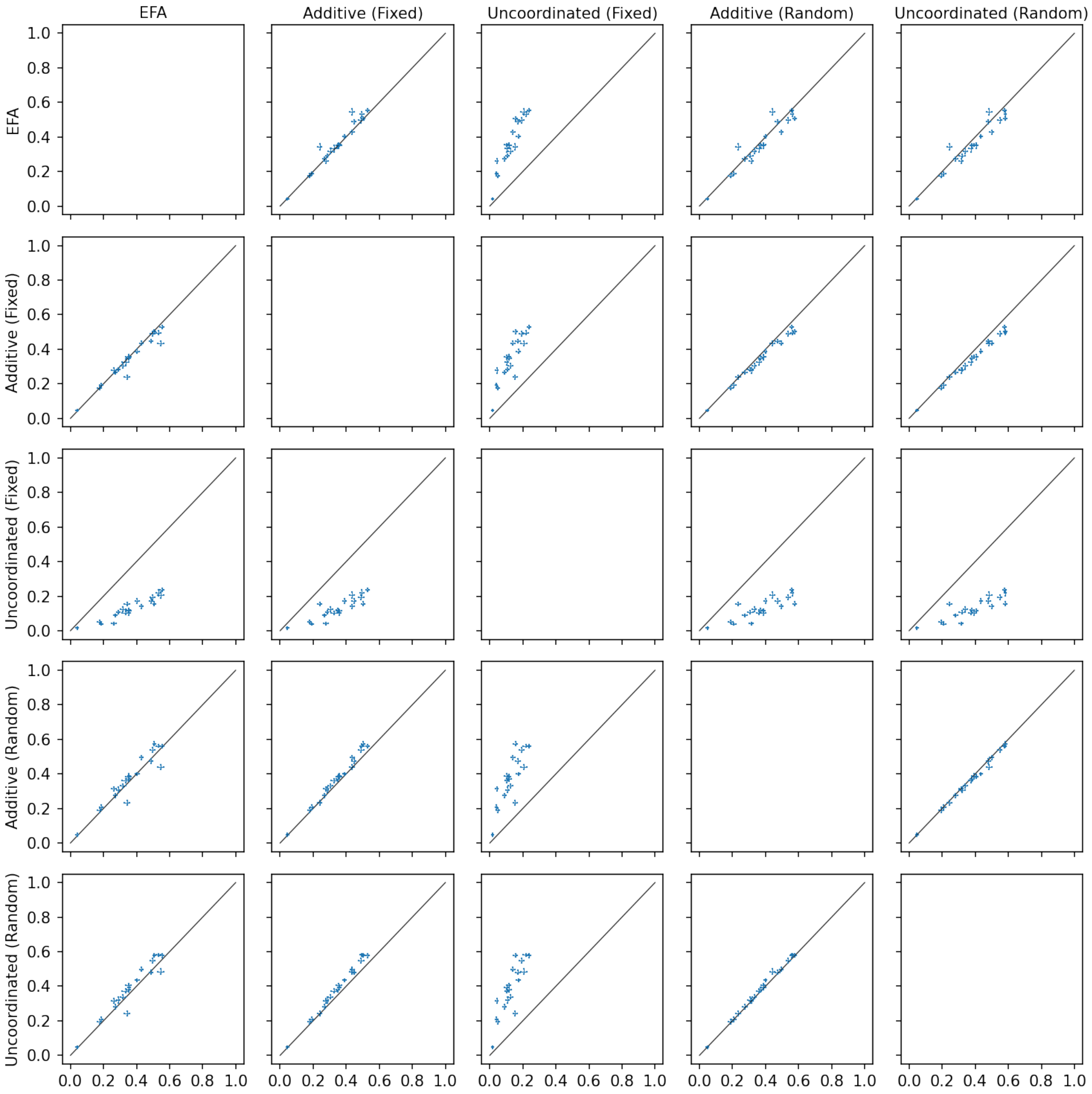

Figure S4: Pairwise comparisons of different polygenic prediction models for the 20 yeast traits in the second dataset. Prediction is measured by the average $R^2$ between the out-of-sample predicted phenotypes and the true phenotypes in 10-fold cross validation.

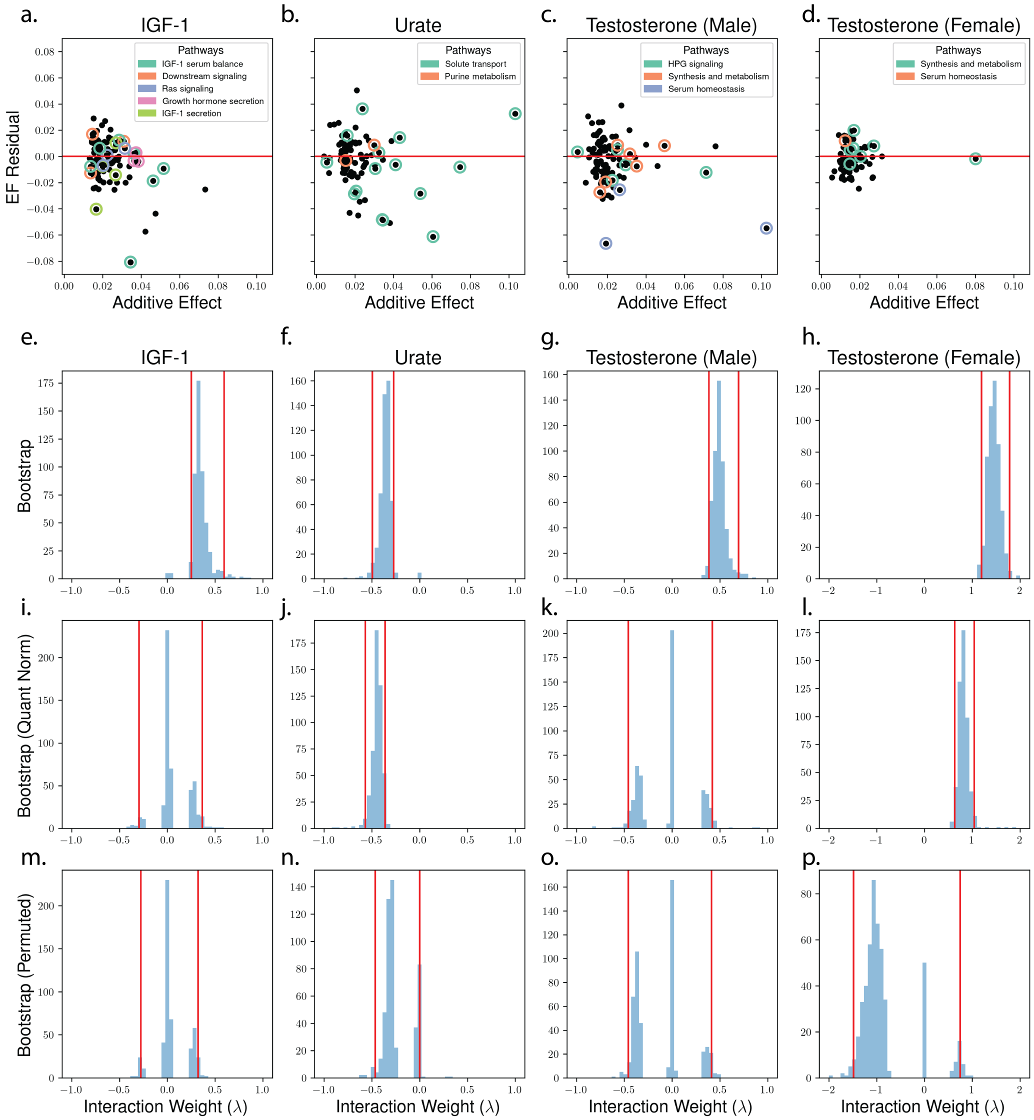

Figure S5: EFA identifies statistically and biologically significant epistasis in 4 human complex traits. (a-d) EFA jointly partitions the additive effects of GWAS-significant SNPs (x-axis) into two EFs (difference in EF weights on the y-axis). (e-f) Bootstrap replicates show that EFA identifies significant epistasis in all four traits. (i-l) Conservatively, we test EFA after quantile-normalizing the phenotypes and EFA remains significant for two traits. (m-p) EFA is not significant for permuted phenotypes. For visualization, 14 total values of $\lambda$ are excluded across all plots (out of a total of 6,000).

The null distribution of $\hat{\lambda}$ is trimodal in (m-p), which is initially surprising. However, this is analogous to well-established bimodal null distributions in linear mixed models [1] (because $\lambda^2$ is bimodal), and is consistent with simulated results under the null shown in **Supplementary Figure S8**.

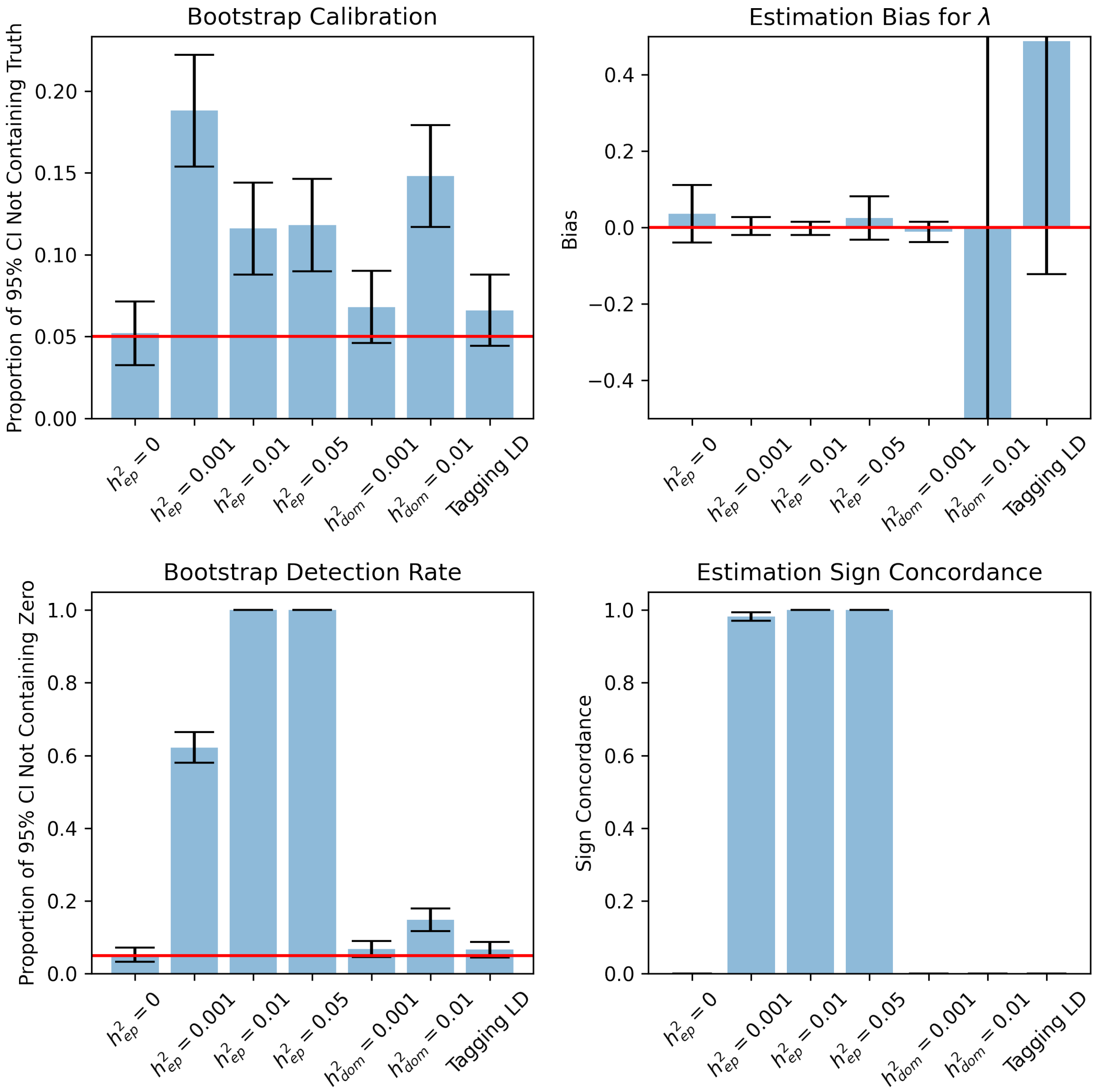

Figure S6: Assessment of the calibration of bootstrap confidence intervals in simulations. All simulations are with with $N = 10000$ individuals, $M = 10$ SNPs, and total heritability $H^2 = 0.1$. Statistics are computed with 500 independent simulations each using 500 bootstrap samples. The parameter $\lambda$ referenced in the estimation bias and sign concordance was fit on the original, un-resampled dataset.

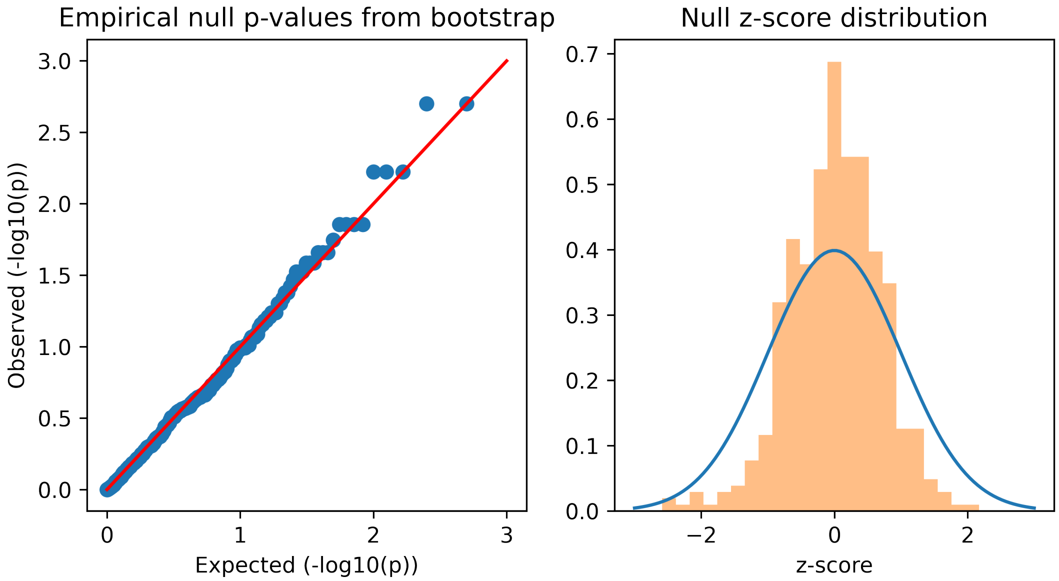

Figure S7: Calibration of p-values obtained from the bootstrap in simulations. Two-sided, empirical p-values calculated as two times the minimum between the fraction of bootstrap samples with $\lambda < 0$ and the fraction of bootstrap samples with $\lambda > 0$. Gaussian approximation z-scores computed as the mean divided by standard deviation.

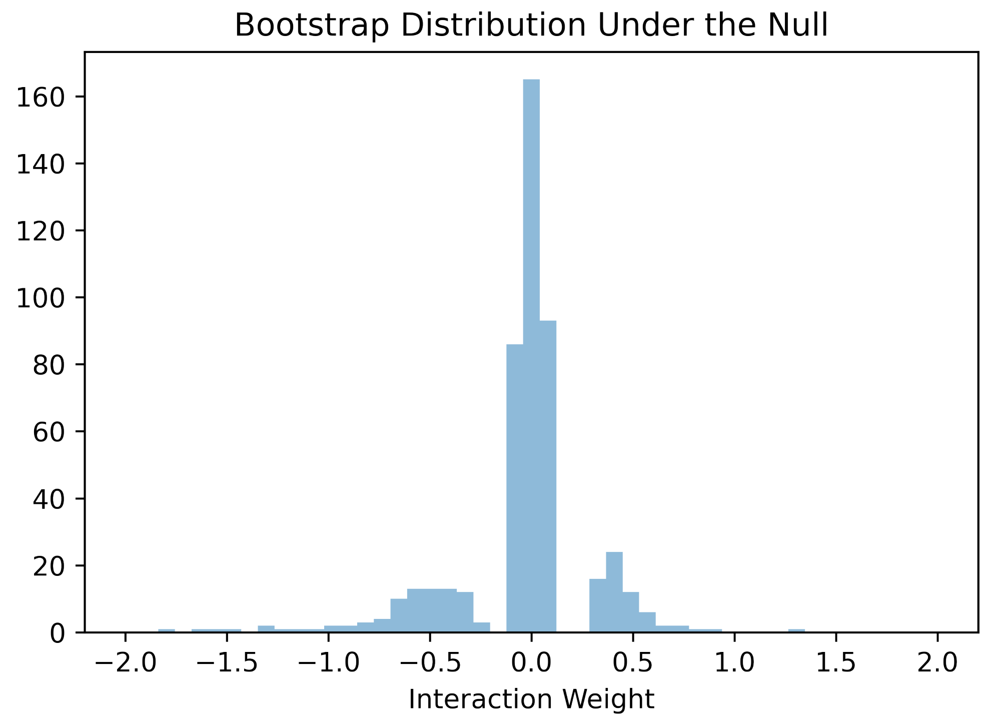

Figure S8: Representative tri-modality of the bootstrap resampling distribution under a null simulation without any epistasis effects. We simulate under a null model with epistasis heritability set to zero and examine the distribution from fitting $\lambda$ on 1000 bootstrap samples using the coordinate descent algorithm with an additive initialization.

### 5 Supplementary Tables

| | EFA | Add | Uncoord | Add (Random) | Uncoord (Random) |
|---|---|---|---|---|---|
| EFA | x | 6 | 46 | 7 | 7 |
| Add | 20 | x | 46 | 3 | 3 |
| Uncoord | 0 | 0 | x | 0 | 0 |
| Add (Random) | 10 | 5 | 44 | x | 2 |
| Uncoord (Random) | 12 | 5 | 44 | 9 | x |

Table S1: Pairwise comparisons between various polygenic prediction models on the first yeast dataset where each entry is read as the number of traits for which the row method beats the column method (paired $t$-test $p < 0.05$)

| | EFA | Add | Uncoord | Add (Random) | Uncoord (Random) |
|---|---|---|---|---|---|
| EFA | x | 7 | 20 | 2 | 2 |
| Add | 3 | x | 20 | 0 | 0 |
| Uncoord | 0 | 0 | x | 0 | 0 |
| Add (Random) | 6 | 6 | 20 | x | 0 |
| Uncoord (Random) | 9 | 15 | 20 | 18 | x |

Table S2: Pairwise comparisons between various polygenic prediction models on the second yeast dataset where each entry is read as the number of traits for which the row method beats the column method (paired $t$-test $p < 0.05$)

| | 10 snps EF1 ($p$) | EF2 ($p$) | 15 snps EF1 ($p$) | EF2 ($p$) | 20 snps EF1 ($p$) | EF2 ($p$) | Window |
|---|---|---|---|---|---|---|---|
| HPG signaling | 0/5 (1.0) | 0/5 (1.0) | 0/5 (1.0) | 0/5 (1.0) | 0/5 (1.0) | 1/5 (0.681) | 0.1mb |
| Serum homeostasis | 0/3 (1.0) | 3/3 (0.001) | 0/3 (1.0) | 3/3 (0.003) | 0/3 (1.0) | 3/3 (0.007) | 0.1mb |
| Synth. & metab. | 0/6 (1.0) | 1/6 (0.478) | 0/6 (1.0) | 2/6 (0.22) | 0/6 (1.0) | 2/6 (0.345) | 0.1mb |
| HPG signaling | 0/8 (1.0) | 1/8 (0.583) | 0/8 (1.0) | 1/8 (0.741) | 0/8 (1.0) | 2/8 (0.503) | 1.0mb |
| Serum homeostasis | 1/5 (0.416) | 3/5 (0.007) | 1/5 (0.564) | 4/5 (0.002) | 1/5 (0.681) | 4/5 (0.005) | 1.0mb |
| Synth. & metab. | 1/7 (0.533) | 1/7 (0.533) | 1/7 (0.692) | 2/7 (0.282) | 1/7 (0.802) | 2/7 (0.426) | 1.0mb |
| HPG signaling | 1/12 (0.739) | 1/12 (0.739) | 2/12 (0.57) | 1/12 (0.875) | 2/12 (0.743) | 2/12 (0.743) | 10.0mb |
| Serum homeostasis | 1/7 (0.533) | 3/7 (0.021) | 1/7 (0.692) | 4/7 (0.009) | 1/7 (0.802) | 4/7 (0.028) | 10.0mb |
| Synth. & metab. | 3/23 (0.415) | 3/23 (0.415) | 3/23 (0.726) | 5/23 (0.236) | 5/23 (0.51) | 6/23 (0.289) | 10.0mb |

Table S3: Biological pathway enrichment in the EFs of male testosterone as a function of # of SNPs assigned to each EF and the window size for assigning SNPs to genes.

| | 10 snps EF1 ($p$) | EF2 ($p$) | 15 snps EF1 ($p$) | EF2 ($p$) | 20 snps EF1 ($p$) | EF2 ($p$) | Window |
|---|---|---|---|---|---|---|---|
| Serum homeostasis | 0/1 (1.0) | 0/1 (1.0) | 1/1 (0.195) | 0/1 (1.0) | 1/1 (0.26) | 0/1 (1.0) | 0.1mb |
| Synth. & metab. | 1/7 (0.638) | 0/7 (1.0) | 2/7 (0.412) | 0/7 (1.0) | 2/7 (0.588) | 0/7 (1.0) | 0.1mb |
| Serum homeostasis | 0/1 (1.0) | 0/1 (1.0) | 1/1 (0.195) | 0/1 (1.0) | 1/1 (0.26) | 0/1 (1.0) | 1.0mb |
| Synth. & metab. | 2/13 (0.536) | 1/13 (0.862) | 3/13 (0.488) | 1/13 (0.955) | 3/13 (0.719) | 1/13 (0.987) | 1.0mb |
| HPG signaling | 0/4 (1.0) | 1/4 (0.434) | 1/4 (0.588) | 1/4 (0.588) | 1/4 (0.708) | 1/4 (0.708) | 10.0mb |
| Serum homeostasis | 0/1 (1.0) | 0/1 (1.0) | 1/1 (0.195) | 0/1 (1.0) | 1/1 (0.26) | 0/1 (1.0) | 10.0mb |
| Synth. & metab. | 5/24 (0.155) | 3/24 (0.662) | 6/24 (0.298) | 3/24 (0.916) | 7/24 (0.434) | 5/24 (0.834) | 10.0mb |

Table S4: Biological pathway enrichment in the EFs of female testosterone as a function of # of SNPs assigned to each EF and the window size for assigning SNPs to genes.

| | 10 snps EF1 ($p$) | EF2 ($p$) | 15 snps EF1 ($p$) | EF2 ($p$) | 20 snps EF1 ($p$) | EF2 ($p$) | Window |
|---|---|---|---|---|---|---|---|
| Downstream signaling | 1/3 (0.284) | 0/3 (1.0) | 1/3 (0.403) | 0/3 (1.0) | 2/3 (0.109) | 0/3 (1.0) | 0.1mb |
| Growth horm. secr. | 0/4 (1.0) | 1/4 (0.361) | 0/4 (1.0) | 1/4 (0.499) | 0/4 (1.0) | 1/4 (0.614) | 0.1mb |
| IGF-1 secretion | 0/4 (1.0) | 1/4 (0.361) | 0/4 (1.0) | 1/4 (0.499) | 0/4 (1.0) | 1/4 (0.614) | 0.1mb |
| IGF-1 serum balance | 0/7 (1.0) | 1/7 (0.549) | 1/7 (0.708) | 1/7 (0.708) | 1/7 (0.817) | 2/7 (0.45) | 0.1mb |
| Ras signaling | 0/5 (1.0) | 1/5 (0.43) | 1/5 (0.581) | 1/5 (0.581) | 1/5 (0.698) | 1/5 (0.698) | 0.1mb |
| Downstream signaling | 1/6 (0.493) | 0/6 (1.0) | 1/6 (0.65) | 0/6 (1.0) | 3/6 (0.103) | 0/6 (1.0) | 1.0mb |
| Growth horm. secr. | 0/6 (1.0) | 2/6 (0.117) | 0/6 (1.0) | 3/6 (0.047) | 0/6 (1.0) | 3/6 (0.103) | 1.0mb |
| IGF-1 secretion | 0/5 (1.0) | 1/5 (0.43) | 0/5 (1.0) | 1/5 (0.581) | 0/5 (1.0) | 1/5 (0.698) | 1.0mb |
| IGF-1 serum balance | 0/9 (1.0) | 2/9 (0.236) | 1/9 (0.799) | 2/9 (0.427) | 1/9 (0.89) | 3/9 (0.279) | 1.0mb |
| Ras signaling | 0/12 (1.0) | 1/12 (0.755) | 1/12 (0.887) | 2/12 (0.596) | 2/12 (0.767) | 2/12 (0.767) | 1.0mb |
| Downstream signaling | 2/12 (0.364) | 0/12 (1.0) | 2/12 (0.596) | 0/12 (1.0) | 4/12 (0.217) | 1/12 (0.95) | 10.0mb |
| Growth horm. secr. | 2/13 (0.406) | 2/13 (0.406) | 2/13 (0.644) | 4/13 (0.117) | 2/13 (0.808) | 4/13 (0.269) | 10.0mb |
| IGF-1 secretion | 0/6 (1.0) | 1/6 (0.493) | 0/6 (1.0) | 1/6 (0.65) | 0/6 (1.0) | 1/6 (0.764) | 10.0mb |
| IGF-1 serum balance | 0/10 (1.0) | 2/10 (0.278) | 1/10 (0.833) | 2/10 (0.487) | 1/10 (0.915) | 3/10 (0.345) | 10.0mb |
| Ras signaling | 4/37 (0.587) | 3/37 (0.822) | 5/37 (0.768) | 6/37 (0.558) | 7/37 (0.731) | 9/37 (0.338) | 10.0mb |

Table S5: Biological pathway enrichment in the EFs of IGF-1 as a function of # of SNPs assigned to each EF and the window size for assigning SNPs to genes.

| | 10 snps EF1 ($p$) | EF2 ($p$) | 15 snps EF1 ($p$) | EF2 ($p$) | 20 snps EF1 ($p$) | EF2 ($p$) | Window |
|---|---|---|---|---|---|---|---|
| Purine metabolism | 0/2 (1.0) | 0/2 (1.0) | 0/2 (1.0) | 0/2 (1.0) | 0/2 (1.0) | 0/2 (1.0) | 0.1mb |
| Solute transport | 3/15 (0.175) | 5/15 (0.007) | 4/15 (0.166) | 6/15 (0.01) | 4/15 (0.355) | 6/15 (0.048) | 0.1mb |
| Purine metabolism | 0/4 (1.0) | 0/4 (1.0) | 0/4 (1.0) | 1/4 (0.487) | 0/4 (1.0) | 1/4 (0.601) | 1.0mb |
| Solute transport | 5/28 (0.111) | 9/28 (< 0.001) | 6/28 (0.213) | 10/28 (0.001) | 6/28 (0.525) | 11/28 (0.005) | 1.0mb |
| Purine metabolism | 0/39 (1.0) | 1/39 (0.995) | 3/39 (0.979) | 3/39 (0.979) | 6/39 (0.89) | 5/39 (0.961) | 10.0mb |
| Solute transport | 8/34 (0.003) | 9/34 (0.0) | 9/34 (0.026) | 11/34 (0.001) | 9/34 (0.194) | 13/34 (0.002) | 10.0mb |

Table S6: Biological pathway enrichment in the EFs of urate as a function of # of SNPs assigned to each EF and the window size for assigning SNPs to genes.
